# P-TEFb promotes cell survival upon p53 activation by suppressing intrinsic apoptosis pathway

**DOI:** 10.1101/2022.05.29.493929

**Authors:** Zhijia Wang, Monika Mačáková, Andrii Bugai, Sergey G. Kuznetsov, Antti Hassinen, Tina Lenasi, Swapnil Potdar, Caroline C. Friedel, Matjaž Barborič

## Abstract

Positive transcription elongation factor b (P-TEFb) is the crucial player in RNA polymerase II (Pol II) pause release that has emerged as a promising target in cancer. Because single-agent therapy may fail to deliver durable clinical response, targeting of P-TEFb shall benefit when deployed as a combination therapy. We screened a comprehensive oncology library and identified clinically relevant antimetabolites and Mouse double minute 2 homolog (MDM2) inhibitors as top compounds eliciting p53-dependent death of colorectal cancer cells in synergy with selective inhibitors of P-TEFb. While the targeting of P-TEFb augments apoptosis by antimetabolite 5-fluorouracil, it switches the fate of cancer cells by the non-genotoxic MDM2 inhibitor Nutlin-3a from cell-cycle arrest to apoptosis. Mechanistically, the fate switching is enabled by the induction of p53-dependent pro-apoptotic genes and repression of P-TEFb-dependent pro-survival genes of the PI3K-AKT signaling cascade, which stimulates Caspase 9 and intrinsic apoptosis pathway in BAX/BAK-dependent manner. Finally, combination treatments trigger apoptosis of cancer cell spheroids. Together, co-targeting of P-TEFb and suppressors of intrinsic apoptosis could become a viable strategy to eliminate cancer cells.

## INTRODUCTION

Gene transcription by RNA polymerase II (Pol II) is a highly regulated process that determines levels of gene expression and, by extension, nearly all facets of cellular identity, fate, and metabolism. A central regulatory hub is the carboxy-terminal domain (CTD) of the largest subunit of Pol II, RPB1, which comprises heptad repeats with the consensus sequence Tyr^1^Ser^2^Pro^3^Thr^4^Ser^5^Pro^6^Ser^7 1^. During the transcription cycle, multiple positions within the CTD repeats are subject to dynamic phosphorylation and dephosphorylation events by transcriptional cyclin-dependent kinases (tCDKs) and phosphatases, respectively. These post-translational modifications modulate the binding of factors required for precursor mRNA (pre-mRNA) initiation, elongation, cotranscriptional maturation, and termination, as well as chromatin modification^2,3^.

Initiation of transcription is stimulated by CDK7, which is together with its regulatory partners cyclin H and Mat1 recruited to the Mediator-Pol II pre-initiation complex on gene promoters to phosphorylate Ser5 and Ser7 residues of the CTD. Concurrently, CDK7 promotes 5′-end capping of transcripts and facilitates promoter-proximal pausing of Pol II by stimulating the recruitment of DRB-sensitivity inducing factor (DSIF) and negative elongation factor (NELF), which bind and arrest Pol II in an inactive conformation^3,4^. Critically, the release of paused Pol II into elongation genome-wide is triggered by the positive transcription elongation factor b (P-TEFb), composed of the catalytic CDK9 and a regulatory cyclin T1 or T2 subunit^5,6^. P-TEFb phosphorylates Ser2 residues across the CTD and its linker, as well as multiple positions within both pause-inducing factors, converting DSIF into a positive elongation factor and triggering dissociation of NELF from Pol II. In turn, the polymerase associated factor complex (PAFc) binds Pol II while SPT6 docks to the CTD linker region, allowing Pol II elongation to resume^7^. Adding another layer of complexity, the P-TEFb-driven release of Pol II into elongation is opposed by phosphoprotein phosphatases, such as a protein phosphatase 2A (PP2A) complex, which is recruited to Pol II by the Integrator complex to dephosphorylate key P-TEFb substrates DSIF and the CTD of Pol II^8-10^. Upon the release of Pol II from pausing, the homologous CDK12 and CDK13 interact with cyclin K to stimulate Pol II elongation throughout gene bodies and facilitate production of full-length transcripts through suppressing their premature cleavage and polyadenylation^11,12^. Finally, while CDK8 and its CDK19 paralogue bind cyclin C to influence gene expression by phosphorylating transcription factors and controlling Mediator structure and function^13^, CDK11 binds cyclins L1 and L2 to regulate transcription and pre-mRNA splicing by phosphorylating the CTD of Pol II and the U2 small nuclear ribonucleoprotein subunit SF3B1^14,15^.

Historically, transcription by Pol II has been deemed too essential to healthy cells for being considered as a target in cancer. Intriguingly, recent work shows that genetic and epigenetic alterations of cancer cells drive a transcriptionally deregulated state, referred to as transcriptional addiction, which facilitates tumor initiation and progression^16,17^. Correspondingly, the addiction, while absent in non-cancerous cells, may instigate a disproportionate reliance on core components of the Pol II machinery to support tumor growth and survival, rendering cancer cells sensitive to perturbation of discrete phases of the transcription cycle^3,17^. Indeed, recent work demonstrated that targeting CDK7, CDK9, and CDK12 with selective and potent small-molecule inhibitors could lead to vigorous anti-tumor effects in xenograft and pre-clinical cancer models^18-21^, conceivably because of heightened demand for specific tCDKs in sustaining oncogene-driven gene expression programs of rapidly dividing cancer cells. Importantly, while effective, the inhibitors of tCDKs imposed minimal toxicity to normal tissues. This is a considerable improvement over preclinical studies and/or clinical trials with pioneering pan-CDK inhibitors such as flavopiridol, which produced undesirable side effects including particularly concerning blood toxicities like neutropenia^22^.

Despite hopeful prospects for targeting tCDKs in cancer, single-agent tCDK interventions may fail to culminate in durable responses due to therapeutic adaptation and/or presence of pre-existing tumor clones refractory to the therapy^23^. Hence, tCDK-centered anti-cancer strategies could benefit when deployed as part of combination therapies. Because combining two compounds could target cancer cells in a synergistic manner, combination therapies enable compounds to be used at lower doses than as single agents, thus offsetting a potential for toxicity and resistance associated with a particular compound. Importantly, while single-agent therapies could lead to cytostatic response, combination approaches might trigger irreversible cell death, a desirable outcome of any effective cancer treatment^24^.

In this study, we began addressing this issue by targeting P-TEFb which has emerged as a promising player in cancer^17,25^. We screened a comprehensive library of oncology compounds to uncover those eliciting cell death in combination with a highly selective P-TEFb inhibitor NVP-2 that effectuates potent anti-cancer activity at non-toxic doses^26,27^. This effort identified several clinically relevant groups of compounds, including top-performing antimetabolites and Mouse double minute 2 homolog (MDM2) inhibitors that activate tumor suppressor p53. By focusing on the pioneering MDM2 inhibitor Nutlin-3a, which dislodges the E3 ubiquitin ligase MDM2 from p53^28^ and thus promises to be a powerful anti-cancer agent for patients with wild-type (WT) p53^29^, we show that selective perturbation of P-TEFb converts the fate of Nutlin-3a-treated cells from cell-cycle arrest to apoptosis. Mechanistically, we uncover that the synthetic lethality of p53 activation and P-TEFb inhibition (PAPI) depends on execution of the intrinsic apoptosis pathway, which is enabled by induction of p53-dependent pro-apoptotic genes and repression of P-TEFb-dependent pro-survival genes encoding components of the PI3K-AKT signaling cascade. Finally, we show that combination treatments within the PAPI framework remain effective in a three-dimensional cell culture setting, suggesting a translational potential of our work.

## RESULTS

### An oncology library screen identifies MDM2 inhibitors and antimetabolites as top compound classes that are synthetic-lethal with selective P-TEFb inhibitor NVP-2

To identify potentially therapeutic compounds that elicit synthetic lethality in combination with highly selective inhibitor of P-TEFb, we conducted an oncology library screen using HCT116 cell line, a well-established model of colorectal cancer encoding the oncogenic KRAS^G13D^ and PIK3CA^H1047R^ proteins (Fig. 1a). We employed FO5A oncology library consisting of 528 investigational and clinical anti-cancer compounds (Supplementary Fig. 1a), and NVP-2, an aminopyrimidine-derived ATP-competitive inhibitor of P-TEFb. We chose NVP-2 over other antagonists of P-TEFb due to its proven antitumor effect in a preclinical setting^26^ as well as its superb potency and selectivity. Namely, NVP-2 inhibits the catalytic activity of P-TEFb and Ser2 phosphorylation of RNA Pol II with half maximal inhibitory concentration [IC_50_] values of 0.5 nM and 4 nM, respectively, and displays at least 1,200-fold lower activity toward CDK1, 2, 4, and 7^27^. To determine the dose of NVP-2 used in the screen, we exposed HCT116 cells to increasing concentrations of NVP-2 and measured cellular viability with luminescence-based assay in which the number of metabolically active cells is determined by quantifying the levels of ATP via luciferase reaction. This approach revealed the IC^50^ value of NVP-2 in HCT116 was 5 nM (Supplementary Fig. 1b). As we aimed to identify compounds eliciting synergistic cell killing in combination with NVP-2, we elected to use 3 nM of NVP-2, which corresponded to its IC30 dose.

**Figure 1.**
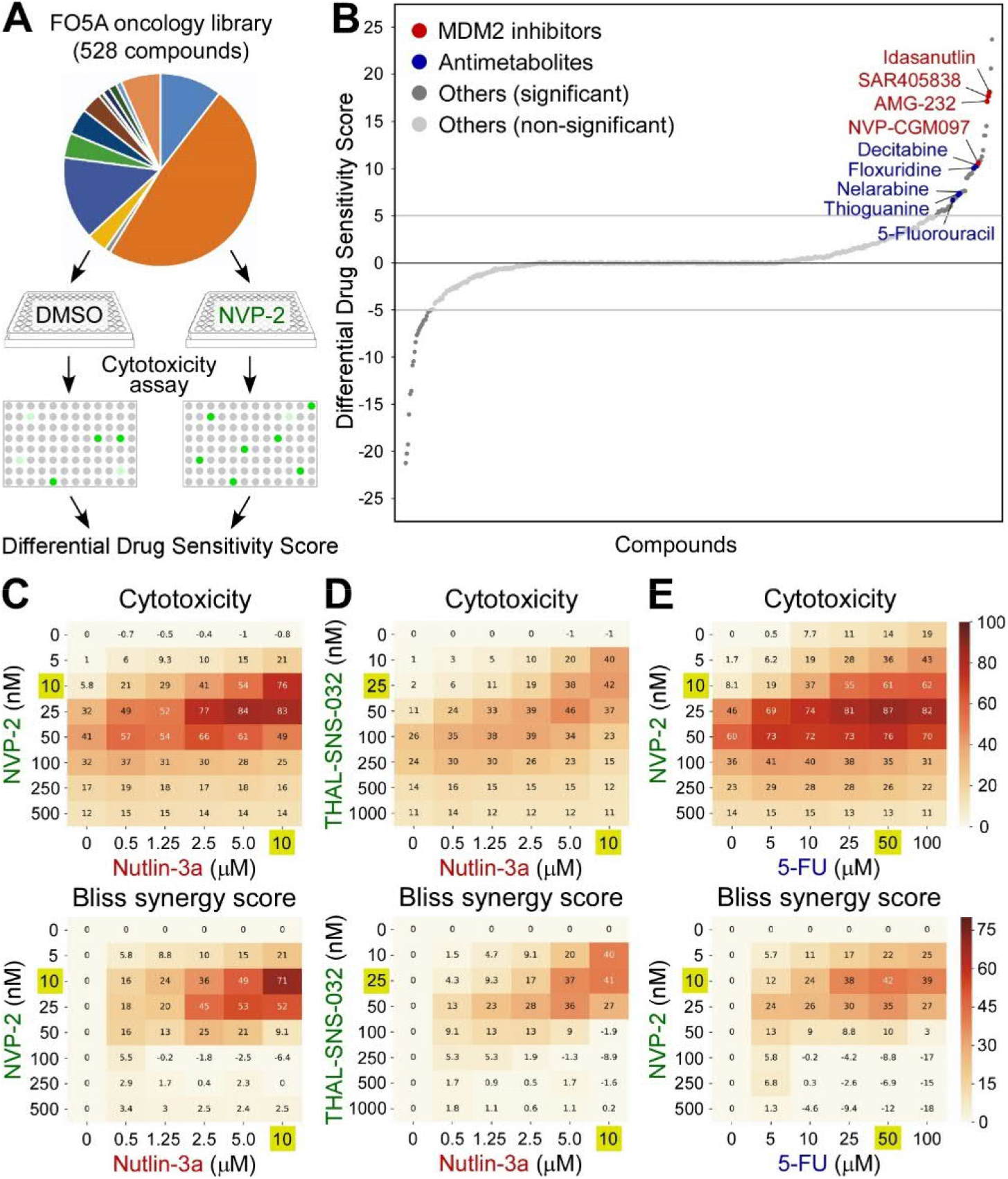
MDM2 inhibitors and antimetabolites are synthetic-lethal with selective perturbation of P-TEFb. (A) Workflow of the FO5A oncology library screen. DMSO- and NVP-2-treated HCT116 cells were exposed to the library compounds at five concentrations for 72 hours, after which cytotoxicity was measured using CellTox Green assay and dDSS values were calculated. Green dots indicate fluorescence readouts of specific FO5A library compound combinations. (B) Waterfall plot representing results of the screen. Compounds with dDSS values ≥ 5 and ≤ -5 were considered as significant (dark grey). MDM2 inhibitors (red) and antimetabolites (blue) are highlighted. (C-E) 8 × 6 cytotoxicity matrices with combinatorial titrations of NVP-2 and THAL-SNS-032 (green) with Nutlin-3a (red) and 5-Fluorouracil (5-FU; blue) at indicated doses to test for the synthetic lethality of compounds in HCT116 cells, depicting cytotoxicity (top) and synergy (bottom) of the combinations. Cytotoxicity values obtained at 48 hr of the treatments using CellTox Green assay were normalized to the DMSO control and are presented as percentages of the maximum cytotoxicity which was set at 100 %. Results represent the average of independent experiments (n = 3). Combinations with the highest Bliss synergy scores are highlighted (gold).

We dispensed HCT116 cells together with DMSO or NVP-2 into 384-well plates containing the FO5A library compounds at five ten-fold serial dilutions. Upon culturing the cells for 72 hours, we measured cellular response to the library compounds using fluorescence-based cytotoxicity assay, in which compromised membrane integrity of dead cells is proportional to fluorescence signal generated by the binding of an otherwise cell impermeable cyanine dye to DNA. To quantify cytotoxicity elicited by the treatments, we generated dose-response curves at each compound concentration and used these to calculate drug sensitivity score (DSS), an area under the dose– response curve type metric as described previously^30^. To identify compounds eliciting cell death in combination with NVP-2, we calculated differential DSS (dDSS) value for each compound by subtracting the DSS values of DMSO-treated cells from NVP-2-treated cells, and considered dDSS values above 5 as effective. This analysis yielded 52 compounds, of which top seven groups include antimetabolites (n = 7), Mouse double minute 2 homolog (MDM2) inhibitors (n = 4), poly(ADP-ribose) polymerase (PARP) inhibitors (n = 4), Inhibitor of apoptosis (IAP) inhibitors/second mitochondria-derived activator of caspase (SMAC) mimetics (n = 3), farnesyl transferase inhibitors (n = 2), Aurora kinase inhibitors (n = 2) and Bromodomain and Extra-Terminal Domain (BET) inhibitor/degrader (n = 2) (Fig. 1b and Supplementary Fig. 1c). These findings indicate that selective targeting of P-TEFb heightens sensitivity of cancer cells to established anti-cancer compounds.

### Amongst transcriptional CDKs, selective targeting of P-TEFb shows the most synergistic lethality with MDM2 inhibitors and antimetabolites

A common denominator of the two leading compound groups of our screen is activation of p53. While the identified antimetabolites are used as chemotherapeutics for a variety of cancers, clinical trials with MDM2 inhibitors have had only a limited success^29^. Nevertheless, the long-standing quest in delivering these inhibitors into the clinic continues, particularly in the context of MDM2 combination therapies. Hence, in this study we investigated how MDM2 inhibitors and antimetabolites elicit synthetic lethality with selective targeting of P-TEFb. As representative compounds, we employed a pioneering MDM2 inhibitor Nutlin-3a and an antimetabolite fluoropyrimidine 5-fluorouracil (5-FU) throughout our work. Whereas Nutlin-3a acts as a competitive protein–protein interaction (PPI) inhibitor of the p53-MDM2 interface by mimicking three hydrophobic amino acid residues of p53 required for MDM2 binding^28^, metabolites of 5-FU interfere with cellular metabolism via incorporating into RNA and DNA and inhibiting the nucleotide synthetic enzyme thymidylate synthase^31^. Even though both compounds activate the p53 pathway in HCT116 cells, non-genotoxic Nutlin-3a triggers reversible cell-cycle arrest in p53-dependent fashion^32^, while genotoxic 5-FU elicits rapid p53-dependent apoptosis^33^.

To confirm the screen results that MDM2 inhibitors and antimetabolites sensitize HCT116 cells to P-TEFb antagonists and to test for synergy of the combination treatments, we first conducted cytotoxicity assays with combinatorial titrations of Nutlin-3a with NVP-2. Whereas Nutlin-3a alone did not show any cytotoxicity across the tested concentrations, it became highly toxic in the presence of sub-lethal doses of NVP-2 (Fig. 1c). Of note, cellular response to NVP-2 alone was biphasic, suggesting that transcriptional shutdown at high NVP-2 doses is incompatible with cell death. We next used Synergyfinder^34^ to determine Bliss synergy scores for all combinations. Here, Bliss score is a readout for synergy depicting the difference between expected effect, which assumes the additive contribution of individual treatments, and observed effect, with positive scores indicating synergy of combination treatment. This analysis showed a striking degree of synergy between high doses of Nutlin-3a and low doses of NVP-2 in inducing lethality of HCT116 cells, wherein the combination treatment of 10 μM of Nutlin-3a with 10 nM of NVP-2 was the most synergistic, reaching the Bliss score of 71 (Fig. 1c, bottom). In contrast, despite an induction of p53 by Nutlin-3a and a decrease in the CTD Ser2 phosphorylation by NVP-2, combinations of Nutlin-3a with NVP2 were not cytotoxic to a non-transformed colorectal epithelial cell line CCD 841 CoN (Supplementary Fig. 1d).

Because NVP-2 can engage CDK10 in addition to P-TEFb at 1 μM dose^27^, we next performed the same cytotoxicity and Bliss synergy score assays, but instead of NVP-2, we employed a selective CDK9 degrader THAL-SNS-032 or a highly selective ATP-competitive P-TEFb inhibitor iCDK9 to perturb the kinase^27,35^. Confirming the results obtained with NVP-2, high doses of Nutlin-3a elicited synthetic lethality with low doses of THAL-SNS-032 and iCDK9 in a synergistic manner, reaching the highest Bliss scores of 41 and 38, respectively (Fig. 1d and Supplementary Fig. 1e). Likewise, increasing doses of 5-FU augmented sensitivity of HCT116 cells to sub-lethal doses of NVP-2, wherein the combination treatment of 50 μM of 5-FU with 10 nM of NVP-2 resulted in the highest Bliss score of 42 (Fig. 1e).

To provide complementary evidence for the synthetic lethality triggered by PAPI, we conducted viability assays with combinatorial titrations of Nutlin-3a and 5-FU with NVP-2, and calculated Bliss synergy scores for all of the combinations. Corroborating the cytotoxicity results, both p53-activating compounds synergized with the sub-lethal doses of NVP-2 to decrease viability of HCT116 cells, reaching the Bliss scores of 49 and 42 with the combination treatments of 10 μM of Nutlin-3a with 25 nm of NVP-2 and 50 μM of 5-FU with 10 nM of NVP-2, respectively (Supplementary Fig. 1f, g). Supporting these findings, 10 μM of Nutlin-3a and 25 μM of 5-FU decreased the IC_50_ value of NVP-2 sharply, which dropped from 5 nM to 1.4 nM and 0.5 nM, respectively (Supplementary Fig. 1b).

Finally, we asked whether specific pharmacological targeting of other major tCDKs can also induce synthetic lethality in combination with Nutlin-3a. We targeted CDK7, CDK8, CDK11 and CDK12/13 using YKL-5-124^36^, Senexin A^37^, OTS964^38^ and THZ531^39^ inhibitors, respectively, and conducted the combinatorial cytotoxicity and Bliss synergy score assays. While targeting CDK8 was not cytotoxic to HCT116 cells, inhibitors of other tCDKs showed modest cytotoxicity in the absence of Nutlin-3a. Importantly, Nutlin-3a treatment elicited robust synthetic lethality only with the inhibitor of CDK7, reaching the highest Bliss score of 44 (Supplementary Fig. 1h-k), which is consistent with a previous report^40^. Together, these findings establish that p53 activating compounds Nutlin-3a and 5-FU elicit cytotoxicity of HCT116 cells in synergy with highly selective antagonists of P-TEFb.

### p53 is required for synthetic lethality of MDM2 inhibitor Nutlin-3a and antimetabolites with P-TEFb inhibitors

Because both Nutlin-3a and 5-FU impact cellular fate via p53, we next evaluated whether the observed synthetic lethality of PAPI depends on p53. To this end, we utilized the parental HCT116 cell line and its isogenic *TP53* ^−/−^ derivative, in which *TP53* was inactivated by disrupting exon 2 of both alleles using recombinant adeno-associated virus (rAAV) vector methodology^41^. We exposed these lines to combinatorial titrations of Nutlin-3a and 5-FU with NVP-2 and compared their cytotoxicity and viability profiles. While the lack of p53 lowered cellular sensitivity to increasing doses of all three compounds administrated alone, it markedly blunted synthetic lethality of both p53 activators with NVP-2 (Fig. 2a-d and Supplementary Fig. 2a-d). Specifically, cytotoxicity of the highly synergistic combinations of Nutlin-3a and 5-FU with NVP-2 dropped in *TP53* null HCT116 cells by 5.7-fold and 7.1-fold, respectively, exemplifying the pivotal contribution of p53 to the synthetic-lethal phenotype (Fig. 2e).

**Figure 2.**
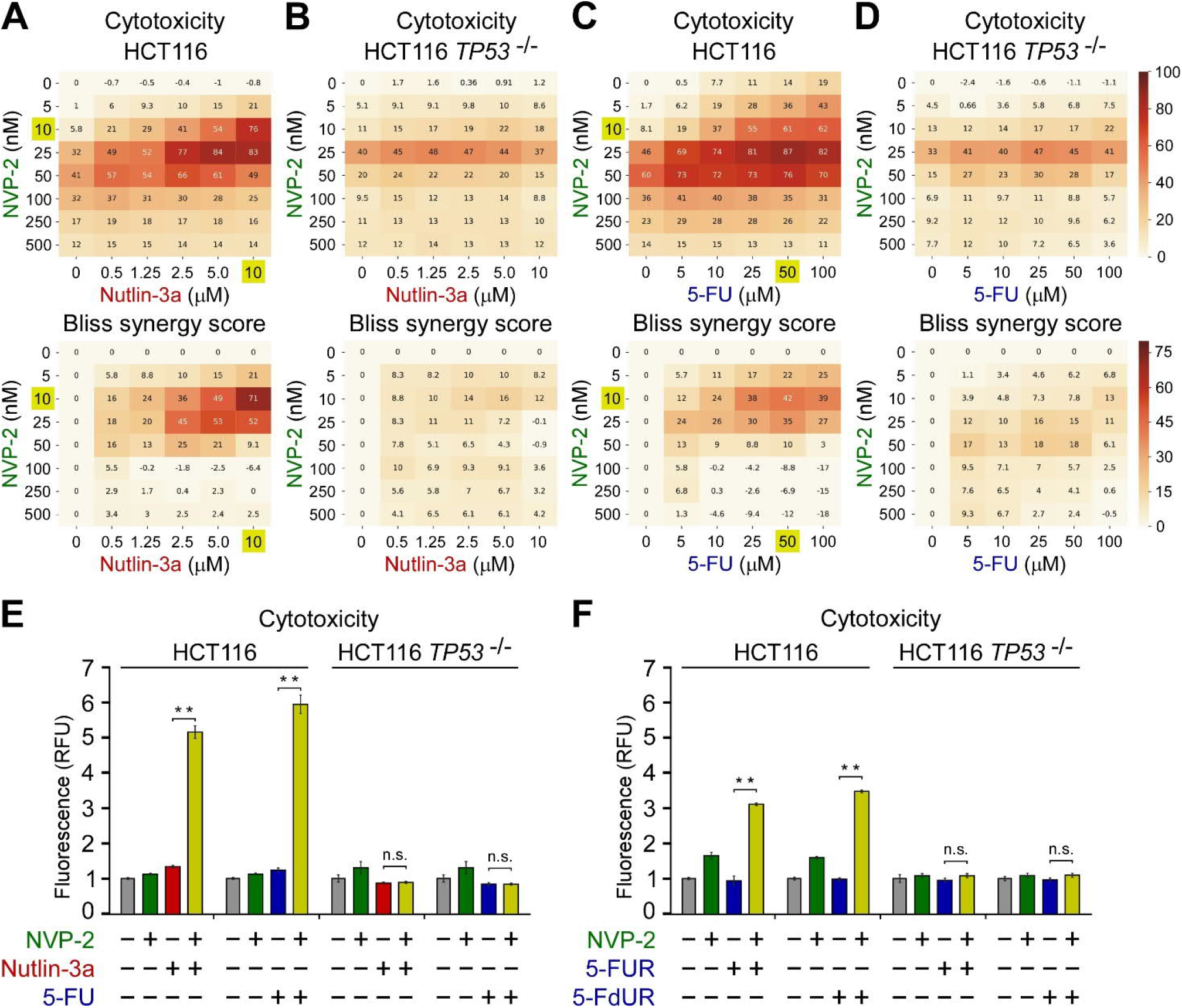
Synthetic lethality of Nutlin-3a and antimetabolites with P-TEFb inhibitor NVP-2 is p53-dependent. (A-D) 8 × 6 matrices with combinatorial titrations of NVP-2 (green) with Nutlin-3a (red) and 5-Fluorouracil (5-FU; blue) at indicated doses to test for the synthetic lethality of compounds in HCT116 and HCT116 *TP53* ^-/-^ cells, depicting cytotoxicity (top) and synergy (bottom) of the combinations. Cytotoxicity values obtained at 48 hr of the treatments using CellTox Green Cytotoxicity Assay were normalized to the values of DMSO-treated cells and are presented as percentages of the maximum cytotoxicity which was set at 100 %. Results represent the average of independent experiments (n = 3). Combinations with the highest Bliss synergy scores in HCT116 cells are highlighted (gold). (E,F) Cytotoxicity of HCT116 and HCT116 *TP53* ^-/-^ cells treated with DMSO (grey), NVP-2 (10 nM; green), Nutlin-3a (10 μM; red), and antimetabolites 5-Fluorouracil (5-FU; 50 μM), 5-fluorouridine (5-FUR; 0.5 μM) and 5-fluorodeoxyuridine (5-FdUR; 0.15 μM) (blue) alone and in combination (gold) as indicated for 48 hr measured using CellTox Green Cytotoxicity Assay. Results are presented as fluorescence values relative to the values of DMSO-treated cells and plotted as the mean ± s.e.m. (n = 3). **, P < 0.01; n.s., non-significant, determined by Student’s *t* test.

Because 5-FU exerts its anti-cancer effects in part through incorporation of its metabolites into RNA and DNA^31^, we next asked whether its synthetic lethality with P-TEFb inhibition depends on RNA or DNA metabolism. To address this question, we utilized ribose and deoxyribose derivatives of 5-FU, 5-fluorouridine (5-FUR) or 5-fluorodeoxyuridine (5-FdUR), respectively. Of note, 5-FdUR, also called Floxuridine, appeared among the top antimetabolites in our oncology library screen (Fig. 1b and Supplementary Fig. 1c) and is used to treat cancer of gastrointestinal tract that has metastasized to the liver. We therefore conducted cytotoxicity assays using the parental and *TP53* null HCT116 cells with combinatorial treatments of 5-FUR and 5-FdUR with NVP-2. We found that toxicity of both 5-FU derivatives was augmented in the presence of NVP-2 in p53-dependent fashion (Fig. 2f), albeit its levels were lower in comparison to 5-FU (Fig. 2e, f). These findings suggest that 5-FU exerts its synthetic lethality with P-TEFb antagonists through RNA and DNA metabolism. Together, p53 enables the synthetic lethality of Nutlin-3a and antimetabolites with P-TEFb antagonists, suggesting that cells become markedly dependent on P-TEFb-driven transcriptional program upon activation of p53.

### p53-dependent activation of apoptosis drives synthetic lethality of p53 activation and P-TEFb perturbation

Upon activation, the DNA-binding transcription factor p53 elicits diverse anti-proliferative transcriptional programs, amongst which cell-cycle arrest and apoptosis are best understood^42^. Based on the evidence gathered thus far, we hypothesized that following non-genotoxic and genotoxic stabilization of p53, perturbation of P-TEFb induces cell death by apoptosis. Therefore, when combined with Nutlin-3a, targeting P-TEFb should convert cell fate choice from cell-cycle arrest to apoptosis, whereas augment apoptosis of cells exposed to sub-lethal doses of 5-FU. To address these predictions, we treated the parental and *TP53* null HCT116 cells with increasing doses of Nutlin-3a and 5-FU in the absence or presence of 10 nM of NVP-2, and conducted immunoblotting analyses to monitor induction of apoptosis and p53 pathway by determining levels of cleaved poly(ADP-ribose) polymerase (cPARP) and p53 along its direct transcriptional target, the CDK inhibitor p21, respectively. Indeed, while Nutlin-3a treatment of the parental HCT116 cells led to stabilization of p53 and induction of p21 in dose-dependent manner, robust cleavage of PARP occurred exclusively in the presence of NVP-2 (Fig. 3a, left). Similarly, the sub-lethal doses of 5-FU activated p53 but augmented cPARP levels only in combination with NVP-2 (Fig. 3b, left). Importantly, none of these occurred in *TP53* null HCT116 cells (Fig. 3a, b, right), once again underscoring the pivotal role of p53 in exerting synthetic lethality of PAPI. Confirming these findings, we observed the same apoptotic phenotype in HCT116 cells treated with Nutlin-3a and 5-FU in combination with selective P-TEFb antagonists THAL-SNS-032 and iCDK9 (Fig. 3c, d and Supplementary Fig. 3a).

**Figure 3.**
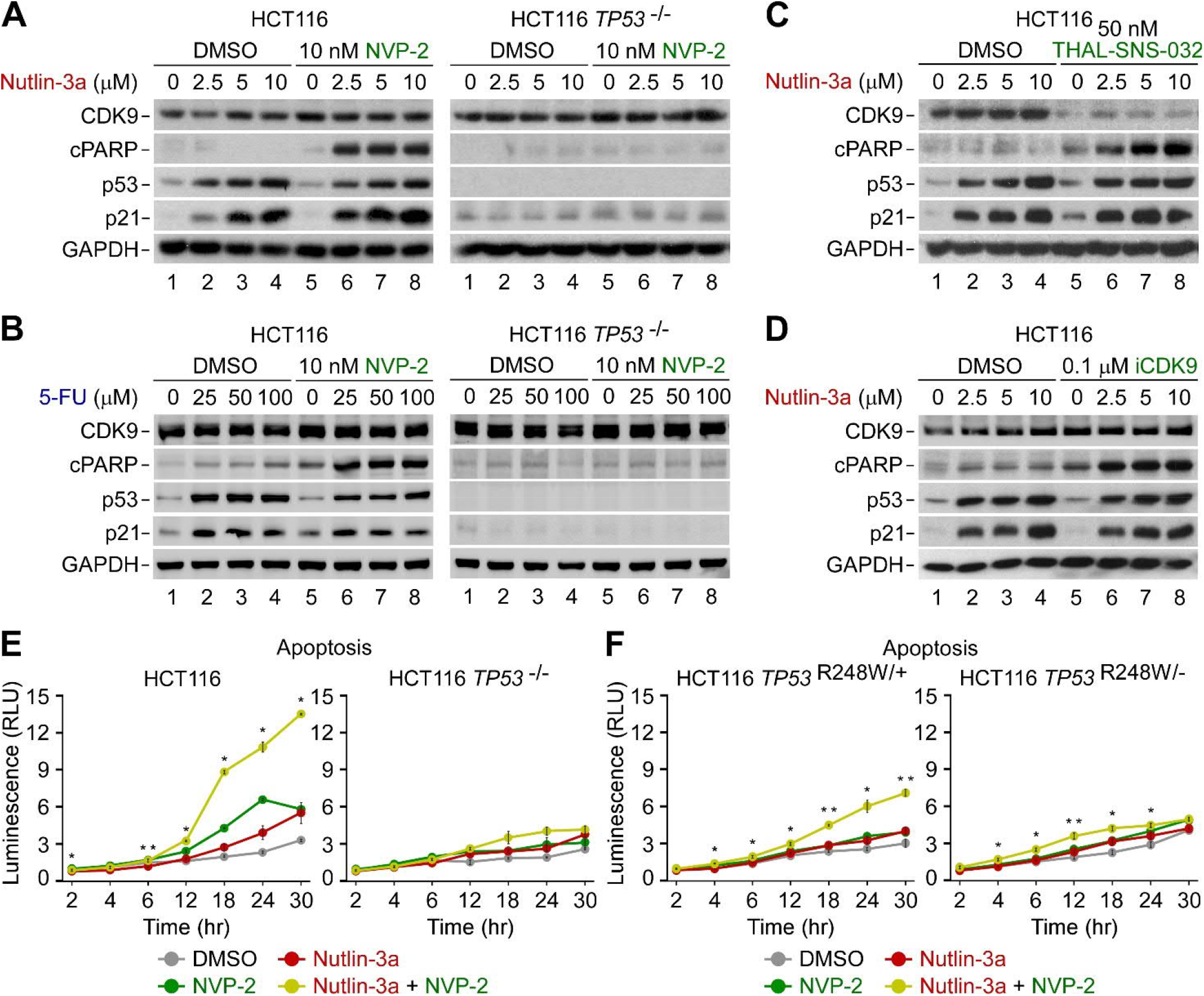
Activation of apoptosis underlies p53-dependent synthetic lethality of p53 activation and P-TEFb inhibition. A-D) HCT116 and HCT116 *TP53* ^-/-^ cells were treated with DMSO and indicated combinations and doses of P-TEFb antagonists (green), Nutlin-3a (red), and 5-Fluorouracil (5-FU; blue) for 24 hr prior to preparation of whole cell extracts and detection of the indicated proteins by Western blotting. (E,F) Apoptosis of the indicated HCT116 cell lines treated with DMSO (grey), NVP-2 (10 nM; green) and Nutlin-3a (10 μM; red) alone and in combination (gold). Results obtained at the time points indicated below the graphs using RealTime-Glo Annexin V Apoptosis and Necrosis Assay are presented as luminescence values relative to the values of DMSO-treated cells at 2 hr and plotted as the mean ± s.e.m. (n = 3). *, P < 0.05; **, P < 0.01, determined by Student’s *t* test using Nutlin-3a and Nutlin-3a + NVP-2 data sets.

To validate these immunoblotting findings and provide a kinetic view of apoptosis activation, we performed a luminescence-based quantitative assay that measures apoptosis-induced exposure of phosphatidylserine on the outer surface of cell membrane through time. The assay uses two Annexin V fusion proteins each containing a fragment of NanoBit Luciferase, of which complementation is achieved upon binding of Annexin V with phosphatidylserine^43^. Compared to low levels of apoptosis elicited by Nutlin-3a and NVP-2 alone, we detected robust and p53-dependent apoptosis of cells exposed to the combination, starting to be apparent at about twelve hours of the treatment (Fig. 3e).

Finally, we asked whether one functional *TP53* allele suffices to support activation of apoptotic cell death following concurrent p53 activation and P-TEFb inhibition. To this end, we performed the immunoblotting and kinetic apoptosis measurements by employing two additional isogenic HCT116 cell line derivatives constructed with the rAAV vector methodology. While the HCT116 *TP53* ^R248W/+^ cell line contains WT and transcriptionally-deficient *TP53* allele harboring R248W mutation found commonly in human cancers, the HCT116 *TP53* ^R248W/-^ cell line lacks functional p53^41^. We treated both cell lines with increasing doses of Nutlin-3a in the absence or presence of 10 nM of NVP-2 and monitored the levels of cPARP, p53 and p21 by immunoblotting. As expected, Nutlin-3a treatment stabilized p53 in both cell lines, while p21 was induced only in HCT116 *TP53* ^R248/+^ cell line, underscoring the transcriptional competence encoded by the WT *TP53* allele (Supplementary Fig. 3b). Critically, Nutlin-3a sensitized HCT116 *TP53* ^R248W/+^ but not HCT116 *TP53* ^R248W/-^ cells to NVP-2-induced apoptosis as documented by increased cleavage of PARP (Supplementary Fig. 3b). Corroborating these findings, kinetic analyses showed robust activation of apoptosis by the combination of Nutlin-3a and NVP-2 over the single-agent treatments only in HCT116 *TP53* ^R248W/+^ cells (Fig. 3f). Of note, the magnitude of apoptosis induced by the combination treatment was approximately two-fold lower compared to the one observed in parental HCT116 cells, whereas the kinetic profiles in both cell lines were very similar (Fig. 3e, f). Together, p53-dependent apoptosis drives synthetic lethality of PAPI, whereby a single WT *TP53* allele is necessary for the effect.

### p53-induced genes of the intrinsic apoptosis pathway remain expressed upon sub-lethal inhibition of P-TEFb

Given the striking ability of P-TEFb antagonists to switch the fate of Nutlin-3a-treated HCT116 cells from undergoing cell-cycle arrest to apoptosis, we next set out to decipher a molecular basis for this phenotype. We reasoned that alterations in transcription could explain the exquisite dependency on P-TEFb following activation of p53. Therefore, we conducted RNA sequencing analysis (RNA-seq) of poly-A enriched pools of RNA isolated from HCT116 cells treated with DMSO, Nutlin-3a and NVP-2 in isolation, and in combination for eight hours, soon after which apoptosis is triggered by the combination treatment (Fig. 3e). Based on the reproducible RNA-seq data (Supplementary Fig. 4a), we determined differentially expressed genes (p-adj ≤ 0.001; Log2 fold-change ≥ 1).

We first analyzed transcriptional changes brought about by Nutlin-3a and examined how selective inhibition of P-TEFb by NVP-2 affected these changes. As expected, Nutlin-3a elicited a highly specific transcriptional program. In comparison to DMSO treatment, expression of only 1.08% of protein-coding genes was altered (n = 10,718), amongst which 112 genes were induced, while four were repressed (Fig. 4a). Using the hallmark gene set collection of the Molecular Signatures Database^44,45^, gene set enrichment analysis revealed that the p53 Hallmark Pathway was the top upregulated gene set within the Nutlin-3a-regulated program (Supplementary Fig. 4b), corroborating the selective targeting of the p53-MDM2 interface by Nutlin-3a. Importantly, the addition of NVP-2 to Nutlin-3a-treated cells significantly impacted expression of the Nutlin-3a-regulated genes (Fig. 4b).

**Figure 4.**
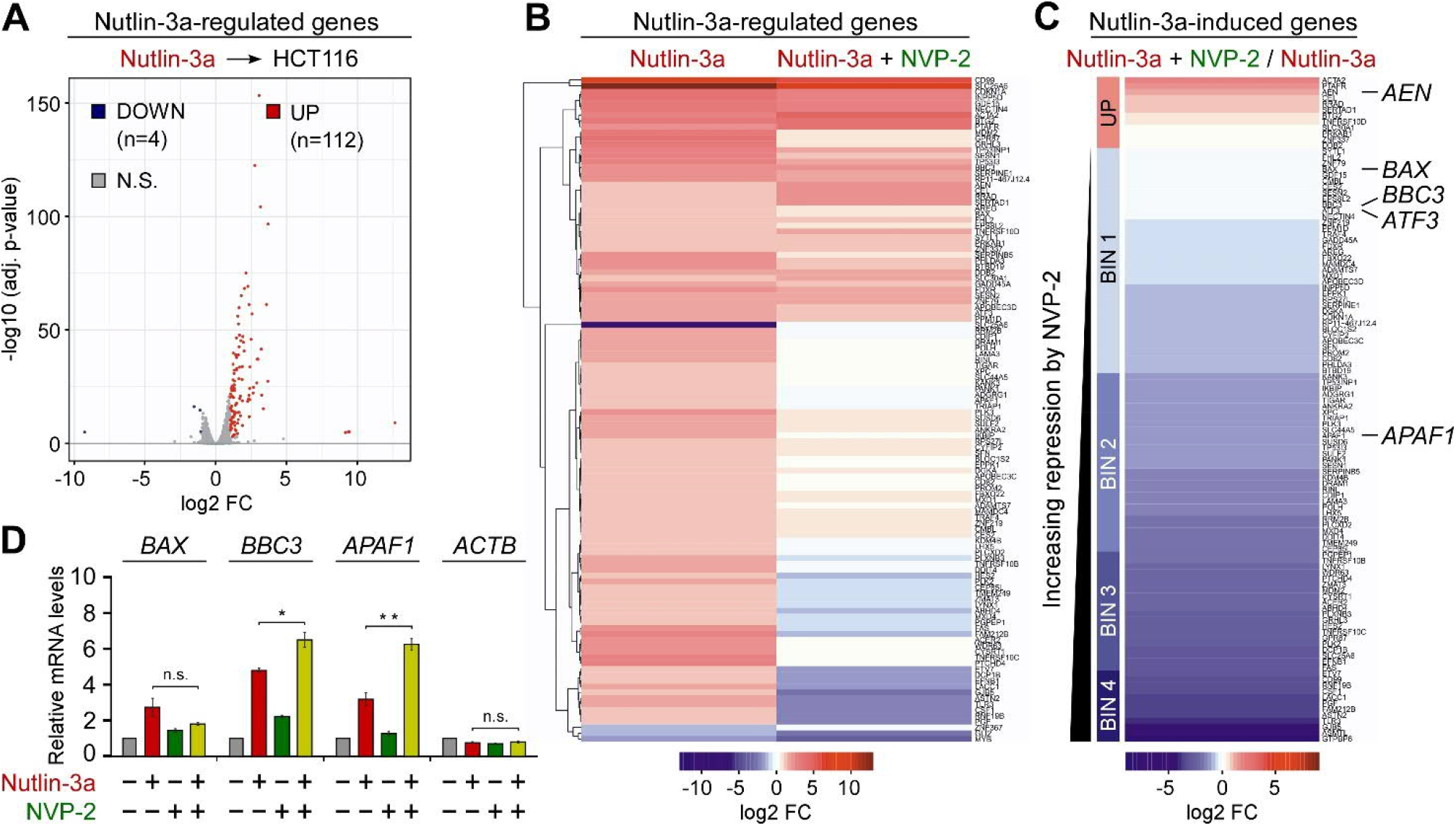
Sub-lethal inhibition of P-TEFb does not block p53-dependent induction of key pro-death genes of the intrinsic apoptosis pathway. (A)Volcano plot representation of differentially expressed protein-coding genes (n = 116; p-adj ≤ 0.001; Log2 FC ≥ 1) from RNA-seq experiments (n = 3) of HCT116 cells treated with DMSO and 10 μM of Nutlin-3a for eight hours. Down-regulated, up-regulated and non-significantly regulated (N.S.) genes are shown as blue, red and grey dots, respectively. FC, fold-change. (B) Hierarchically clustered heatmap representation of Nutlin-3a-regulated protein-coding genes (n = 116; p-adj ≤ 0.001; Log2 FC ≥ 1) from RNA-seq experiments (n = 3) of HCT116 cells treated with 10 μM of Nutlin-3a alone and in combination with 20 nM of NVP-2 for eight hours. Rows represent Log2 FC values of each gene under the depicted treatments on top relative to the values of DMSO-treated cells. FC, fold-change. (C) Heatmap representation of Nutlin-3a-induced protein-coding genes (n = 112; p-adj ≤ 0.001; Log2 FC ≥ 1) from RNA-seq experiments (n = 3) in HCT116 cells treated with 10 μM of Nutlin-3a alone and in combination with 20 nM of NVP-2 for eight hours. Rows represent Log2 FC values of each gene under the Nutlin-3a + NVP-2 combination treatment relative to the values of Nutlin-3a-treated cells. Genes are sorted from largest to smallest Log2 FC values. UP labels genes induced by NVP-2 (Log2 FC >0). BIN 1 (0 > Log2 FC >= -1), BIN 2 (−1 > Log2 FC >= -2), BIN 3 (−2 > Log2 FC >= -3) and BIN 4 (Log2 FC <-3) labels genes repressed by NVP-2. FC, fold-change. (D) HCT116 cells were treated with DMSO (grey), Nutlin-3a (10 μM; red) and NVP-2 (10 nM; green) alone and in combination (gold) as indicated for 24 hr prior to quantifying mRNA levels of the indicated genes with RT-qPCR. Results normalized to the levels of *GAPDH* mRNA and DMSO-treated cells are presented as the mean ± s.e.m. (n = 3). *, P < 0.05; **, P < 0.01; n.s., non-significant, determined by Student’s *t* test.

Because transcriptional activation by p53 is critical for triggering apoptosis^46^, we next focused on the 112 set of genes induced by Nutlin-3a and quantified the impact of P-TEFb inhibition on this set of p53 targets. To this end, we calculated log2 fold-changes in expression for these genes in cells co-treated with Nutlin-3a and NVP-2 compared to Nutlin-3a treatment alone and created a heatmap with genes rank-ordered by decreasing changes in expression (Fig. 4c). This analysis revealed that expression of twelve Nutlin-3a-induced genes increased further upon the addition of NVP-2, while expression of the remaining 100 genes either did not increase as much as with Nutlin-3a treatment alone (35 genes with log2 fold-change ≥ 1 compared to DMSO, 77 with log2 fold-change > 0) or decreased in expression compared to pre-Nutlin-3a levels (33 with log2 fold-change < 0 compared to DMSO). We next divided these 100 genes into four bins based on the magnitude of repression by NVP-2. Given that Nutlin-3a synergizes with NVP-2 to induce cell death by apoptosis, we reasoned that genes critical to the onset of apoptosis should not be exceedingly repressed by NVP-2. Indeed, we identified pro-apoptotic genes of the mitochondria-mediated intrinsic apoptosis pathway, including *AEN, BAX, BBC3* and *ATF3*, amongst genes showing no or little repression by NVP-2 (BIN 1), while another key member of this pathway, *APAF1*, was repressed only moderately (BIN 2).

*BAX, BBC3*, and *APAF1*, which code for BCL-2-associated X protein (BAX), p53-upregulated modulator of apoptosis (PUMA) and apoptotic protease-activating factor 1 (APAF1), respectively, are well-established regulators of the intrinsic apoptosis pathway^47^. Here, the balance between pro-apoptotic and pro-survival members of the BCL-2 family of proteins governs mitochondrial outer membrane permeabilization (MOMP), the key commitment step to undergo apoptotic cell death^48^. Specifically, pro-apoptotic members, such as BCL-2-interacting mediator of cell death (BIM), truncated BH3-interacting domain death agonist (tBID) and PUMA, can be bound and sequestered by pro-survival members such as BCL-2, B cell lymphoma extra large (BCL-X_L_) or myeloid cell leukaemia 1 (MCL1)^49^. However, when these pro-survival proteins are saturated or absent, the balance is tipped in favor of the pro-apoptotic members, enabling MOMP by the pore-forming BAX and BCL-2 antagonist/killer (BAK) oligomers, which releases apoptogenic molecules such as cytochrome *c* from the mitochondrial intermembrane space. In turn, cytochrome *c* binds APAF1 in the cytosol to assemble the heptameric apoptosome^50^ that serves as a platform for autocleavage of the initiator procaspase 9, of which the active form leaves apoptosome to stimulate the downstream executioner caspases 3 and 7, culminating in dismantling of the cell.

Therefore, we next independently assessed our RNA-seq findings by conducting RT-qPCR analysis to determine mRNA levels of *BAX, BBC3*, and *APAF1*. Because 10 μM of Nutlin-3a exhibited the highest synergy in eliciting synthetic lethality with 10 nM of NVP-2 (Fig. 1c), we exposed HCT116 cells to these treatments individually and in combination. Supporting the RNA-seq data, we observed that Nutlin-3a-induced mRNA levels of *BAX* were decreased only moderately by NVP-2, while those of *BBC3* and *APAF1* were elevated even further (Fig. 4d). Together, these findings show that inhibiting P-TEFb with sub-lethal doses of NVP-2 leaves transcription of p53-dependent genes critical for the intrinsic apoptosis pathway largely unhindered.

### Intrinsic apoptosis pathway drives synthetic lethality of non-genotoxic p53 activation and P-TEFb inhibition in transcription-dependent manner

The above gene expression findings raised two inviting predictions. First, the synthetic lethality of PAPI by Nutlin-3a and NVP-2 could be driven by activation of the intrinsic apoptosis pathway. Second, an efficient blockade of Pol II transcription should attenuate synthetic lethality of the combination treatment as induction of p53-dependent targets including key pro-apoptotic genes would not occur.

To address the requirement of the intrinsic apoptosis pathway for the synthetic lethality, we used the parental HCT116 cell line and its isogenic *BAX*/*BAK1* ^−/−^ derivative that lacks the critical mediators of MOMP, treated each cell line with 10 μM of Nutlin-3a and 10 nM of NVP-2 alone and in combination, and performed cytotoxicity assay. Strikingly, in contrast to the parental HCT116 cells, the *BAX*/*BAK1* null cells were completely resistant to cell death elicited by the combination treatment (Fig. 5a). Furthermore, we monitored activity of the initiator caspase 9 in cell extracts of both cell lines exposed to the treatments by utilizing a luminescence-based quantitative assay that measures cleavage of a pro-luminogenic substrate by caspase 9. Critically, the combination treatment led to synergistic activation of caspase 9 in the parental cells, whereas the *BAX*/*BAK1* null HCT116 cells showed no activation (Fig. 5b). Thus, the synthetic lethality of non-genotoxic p53 activation and P-TEFb inhibition depends on the intrinsic apoptosis pathway.

**Figure 5.**
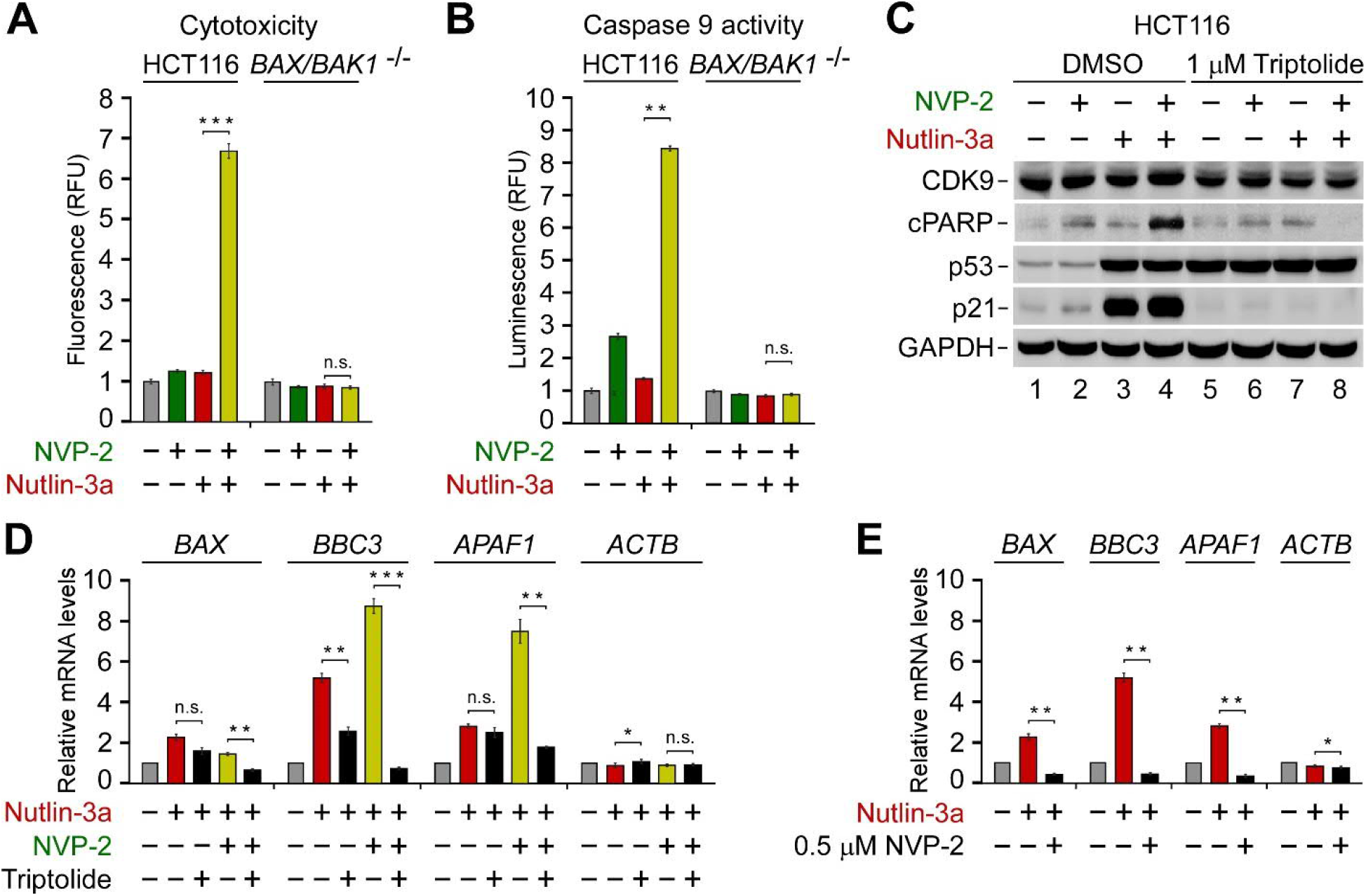
Synthetic lethality of non-genotoxic p53 activation and P-TEFb inhibition depends on intrinsic apoptosis pathway and on-going Pol II transcription. (A) Cytotoxicity of HCT116 and HCT116 *BAX*/*BAK1* ^-/-^ cells treated with DMSO (grey), NVP-2 (10 nM; green) and Nutlin-3a (10 μM; red) alone and in combination (gold) as indicated for 48 hr measured using CellTox Green Cytotoxicity Assay. Results are presented as fluorescence values relative to the values of DMSO-treated cells and plotted as the mean ± s.e.m. (n = 3). ***, P < 0.001; n.s., non-significant, determined by Student’s *t* test. (B) Activity of Caspase 9 measured using Caspase-Glo 9 Assay in whole cell extracts of HCT116 and HCT116 *BAX*/*BAK1* ^-/-^ cells treated with DMSO (grey), NVP-2 (3 nM; green) and Nutlin-3a (10 μM; red) alone and in combination (gold) as indicated for 18 hr. Results are presented as luminescence values relative to the values of DMSO-treated cells and plotted as the mean ± s.e.m. (n = 4). **, P < 0.01; n.s., non-significant, determined by Student’s *t* test. (C) HCT116 cells treated with indicated combinations of NVP-2 (10 nM; green) and Nutlin-3a (10 μM; red) were co-treated with DMSO and Triptolide as indicated for 24 hr prior to preparation of whole cell extracts and detection of the indicated proteins by Western blotting. (D) HCT116 cells were treated with DMSO (grey), Nutlin-3a alone (10 μM; red) and in combination with NVP-2 (10 nM; gold), and Triptolide (1 μM; black) as indicated for 24 hr prior to quantifying mRNA levels of the indicated genes with RT-qPCR. Results normalized to the levels of *GAPDH* mRNA and DMSO-treated cells are presented as the mean ± s.e.m. (n = 3). *, P < 0.05; **, P < 0.01; ***, P < 0.001; n.s., non-significant, determined by Student’s *t* test. (E) HCT116 cells were treated with DMSO (grey), Nutlin-3a (10 μM; red) and NVP-2 (0.5 μM; black) as indicated for 24 hr prior to quantifying mRNA levels of the indicated genes with RT-qPCR. Results normalized to the levels of *GAPDH* mRNA and DMSO-treated cells are presented as the mean ± s.e.m. (n = 3). *, P < 0.05; **, P < 0.01, determined by Student’s *t* test.

To determine whether the synthetic lethal phenotype requires Pol II transcription, we first employed triptolide which inhibits Pol II transcription by blocking the ATPase activity of the TFIIH subunit XPB^51^. Consistent with our prediction, triptolide blunted cell death of HCT116 cells exposed to the combinatorial titrations of Nutlin-3a with NVP-2 (Supplementary Fig. 5a). To gather complementary evidence for this phenotype, we treated HCT116 cells with 10 μM of Nutlin-3a and 10 nM of NVP-2 alone and together in the absence or presence of triptolide and monitored the levels of CDK9, cPARP, p53 and p21 by immunoblotting. While triptolide stabilized p53 across all conditions, it abrogated cleavage of PARP and induction of p21 (Fig. 5c), confirming that blocking Pol II transcription is incompatible with apoptosis triggered by the combination treatment. Likewise, triptolide prevented activation of caspase 9 in cells co-treated with Nutlin-3a and NVP-2 (Supplementary Fig. 5b). Importantly, we monitored mRNA levels of the intrinsic apoptosis genes *BAX, BBC3*, and *APAF1* under the same treatments using RT-qPCR. Consistent with our cytotoxicity and apoptosis findings, triptolide ablated induction of *BAX, BBC3*, and *APAF1* under the combination treatment (Fig. 5d), the effect we recapitulated by substituting triptolide with the 500 nM dose of NVP-2 (Fig. 5e). These findings are consistent with our initial observation that increasing the dose of P-TEFb antagonists beyond the one eliciting the maximal cytotoxicity precluded cell death when administered alone and in combination with Nutlin-3a or 5-FU (Fig. 1 and 2). Corroborating this notion, the levels of cPARP and the p53-induced p21 protein peaked at 50 nM of NVP-2 and 500 nM of iCDK9, while higher doses of both inhibitors diminished these markers of apoptosis and p53-dependent transcription (Supplementary Fig. 5c, d). Together, stimulation of the intrinsic apoptosis pathway drives the switch of Nutlin-3a-treated HCT116 cells from cell-cycle arrest to apoptosis, wherein p53-dependent induction of key pro-apoptotic genes plays a critical role.

### Inhibition of P-TEFb hampers PI3K-AKT pathway via transcriptional repression of key components of PI3K signaling

While our findings thus far showed that the sub-lethal dose of NVP-2 did not impede a key element of the pro-apoptotic transcriptional program by p53, it remained unclear why do cells become markedly dependent on P-TEFb upon activation of p53. To this end, we next focused on transcriptional changes elicited by NVP-2. Compared to Nutlin-3a, NVP-2 treatment had a much broader impact on gene expression, underscoring a pivotal role of P-TEFb in transcription by Pol II. Applying our criteria, we identified 22.69% differentially expressed protein-coding genes (n = 10,728), amongst which 2137 genes were repressed, while 297 were induced (Fig. 6a). These widespread alterations in transcription very likely affect a variety of processes, many of which might promote apoptosis of HCT116 cells treated by Nutlin-3a. Nevertheless, we suspected that a key oncogenic pathway might get impacted by inhibition of P-TEFb, enabling apoptosis of the combination treatment.

**Figure 6.**
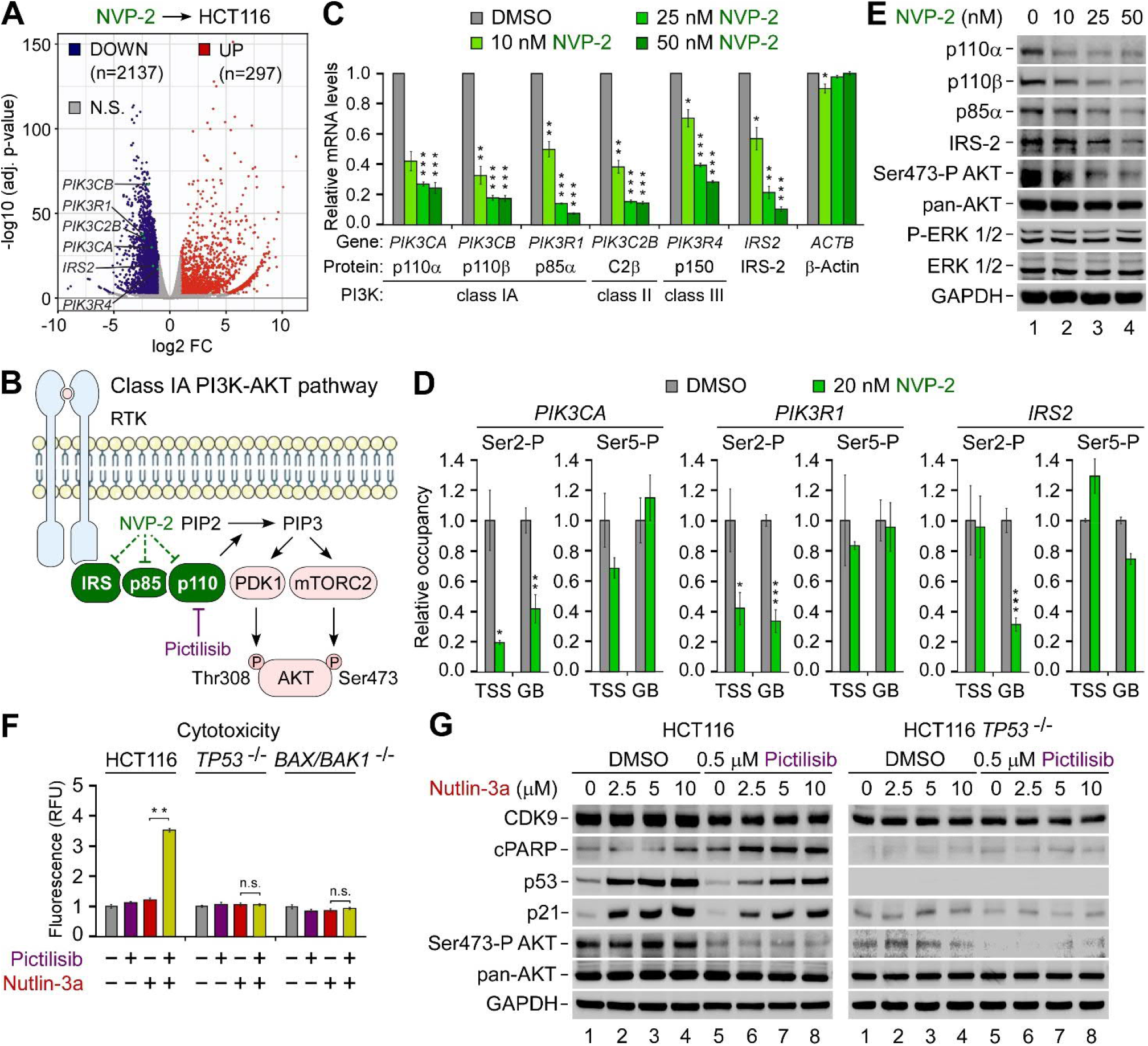
Repression of P-TEFb-dependent genes encoding key components of the pro-survival PI3K-AKT pathway is a driver of the synthetic lethality of p53 activation and P-TEFb inhibition. (A) Volcano plot representation of differentially expressed protein-coding genes (p-adj ≤ 0.001; Log2 FC ≥ 1) from RNA-seq experiments (n = 3) of HCT116 cells treated with DMSO and 20 nM of NVP-2 for eight hours. Down-regulated (DOWN), up-regulated (UP) and non-significantly regulated (N.S.) genes are shown as blue, red and grey dots, respectively. FC, fold-change. (B) Cartoon depicting Class IA PI3K-AKT pathway. Ligand-induced activation of transmembrane receptor tyrosine kinase (RTK; blue) results in autophosphorylation on tyrosine residues, recruiting the PI3K catalytic subunit p110α/β/δ to the membrane via PI3K regulatory subunits, such as p85α, of which SH2 domains bind phosphotyrosyl residues on RTKs or adaptor proteins, e.g. IRS-2. Allosterically activated, PI3K phosphorylates phosphatidylinositol 4,5-bisphosphate (PtdIns(4,5)P_2_; PIP2) to produce phosphatidylinositol 3,4,5-trisphosphate (PtdIns(3,4,5)P_3_; PIP3). PIP3 acts as a second messenger to recruit the serine/threonine protein kinase AKT to the plasma membrane alongside phosphoinositide-dependent kinase-1 (PDK-1) and mechanistic target of rapamycin complex 2 (mTORC2), which activate AKT fully by phosphorylating Thr308 and Ser473, respectively. Green dashed lines denote transcriptional repression of genes encoding IRS-2, p85α and p110 by NVP-2. Pharmacological targeting of p110α/β/δ by the class Ia PI3K inhibitor Pictilisib is in magenta. (C) HCT116 cells were treated with DMSO (grey) and increasing doses of NVP-2 (green) for 8 hr prior to quantifying mRNA levels of the indicated genes with RT-qPCR. Proteins encoded by the genes and their PI3K protein family classes are indicated. Results normalized to the levels of *GAPDH* mRNA and DMSO-treated cells are presented as the mean ± s.e.m. (n = 3). *, P < 0.05; **, P < 0.01; ***, P < 0.001, determined by Student’s *t* test. (D) HCT116 cells were treated with DMSO (grey) and NVP-2 (20 nM; green) for 3 hr prior to determining the levels of Ser2-P and Ser5-P forms relative to total Pol II at transcription start site (TSS) and gene body (GB) of the indicated genes with ChIP-qPCR. Results normalized to the values of DMSO-treated cells that were set to 1 are presented as the mean ± s.e.m. (n = 2). *, P < 0.05; **, P < 0.01; ***, P < 0.001, determined by Student’s *t* test. (E) HCT116 cells were treated with DMSO and increasing doses of NVP-2 (green) for 24 hr prior to preparation of whole cell extracts and detection of the indicated proteins by Western blotting. (F) Cytotoxicity of HCT116, HCT116 *TP53* ^-/-^, and HCT116 *BAX*/*BAK1* ^-/-^ cells treated with DMSO (grey), Pictilisib (1 μM; magenta), and Nutlin-3a (10 μM; red) alone and in combination (gold) as indicated for 48 hr measured using CellTox Green Cytotoxicity Assay. Results are presented as fluorescence values relative to the values of DMSO-treated cells and plotted as the mean ± s.e.m. (n = 3). **, P < 0.01; n.s., non-significant, determined by Student’s *t* test. (G) HCT116 and HCT116 *TP53* ^-/-^ cells were treated with DMSO and indicated combinations and doses of Pictilisib (magenta) and Nutlin-3a (red) for 24 hr prior to preparation of whole cell extracts and detection of the indicated proteins by Western blotting.

Amongst the set of genes repressed by NVP-2, we were very intrigued to find genes encoding key components of the oncogenic phosphoinositide 3-kinase (PI3K)–AKT signaling pathway, including class IA PI3K catalytic subunits p110α (*PIK3CA*) and p110β (*PIK3CB*), their regulatory subunit p85α (*PIK3R1*), class II PI3K catalytic subunit C2β (*PIK3C2B*), class III PI3K regulatory subunit p150 (*PIK3R4*), and a receptor tyrosine kinase (RTK) adaptor protein IRS-2 (*IRS2*) (Fig. 6b). PI3K-AKT is the most commonly deregulated pathway in human cancers^52^. Analyses across cancer lineages identified *PIK3CA a*s the most frequently mutated single oncogene^53^, of which the H1047R hotspot mutation within the kinase domain of p110α, also present in HCT116 cells, circumvents the activation route of WT PI3K by promoting localization of the mutant kinase to plasma membrane^54^. There, PI3K phosphorylates phosphatidylinositol 4,5-bisphosphate (PtdIns(4,5)P_2_) to produce the second messenger phosphatidylinositol 3,4,5-trisphosphate (PtdIns(3,4,5)P_3_), leading to the recruitment and full activation of the serine/threonine protein kinase AKT, promoting cell survival, growth and proliferation (Fig. 6b). Importantly, AKT antagonizes intrinsic apoptosis pathway by phosphorylation of pro-apoptotic mediators BAD and caspase 9^55^.

We hypothesized that selective inhibition of P-TEFb interferes with the signaling through the PI3K-AKT pathway at the level of target gene transcription, thus contributing to apoptosis of Nutlin-3a-treated HCT116 cells. We first conducted RT-qPCR analysis using HCT116 cells treated with increasing doses of NVP-2 to assess mRNA levels of *PIK3CA, PIK3CB, PIK3R1, PIK3C2B, PIK3R4* and *IRS2*. We found that their levels decreased in a dose-dependent manner while mRNA levels of *ACTB* remained largely unaffected, which validated the RNA-seq findings (Fig. 6c). To provide further evidence that transcription of the PI3K-related genes depends on P-TEFb, we next employed quantitative chromatin immunoprecipitation (ChIP-qPCR) assays to monitor the occupancy of total Pol II as well as its CTD Ser2 and Ser5 phosphorylated forms at the transcription start site (TSS) and within gene body (GB) of three PI3K genes critical for the signaling through class IA PI3K using DMSO- and NVP-2-treated HCT116 cells. Namely, while CDK7 regulates Pol II transcription by phosphorylating Ser5 and Ser7 residues of the CTD^3^, P-TEFb does so by targeting Ser2 residues^5,6^. To account for the changes in Pol II occupancy at the genes elicited by NVP-2, we normalized signals of the Ser2 and Ser5 phosphorylated forms to those of total Pol II. This analysis showed that a sub-lethal dose of NVP-2 resulted in significant drop of the CTD Ser2 but not Ser5 levels at most tested locations of *PIK3CA, PIK3R1* and *IRS2* (Fig. 6d), confirming that NVP-2 decreases transcription of this set of PI3K-AKTpathway genes by targeting P-TEFb.

On the basis of these results, we reasoned that inhibition of P-TEFb should decrease protein levels of class IA PI3K heterodimers and adaptor protein IRS-2, leading to lowered activity of the PI3K-AKT signaling (Fig. 6e). Because mTORC2 activates AKT by phosphorylating Ser473 (Ser473-P AKT)^52^, we used phospho-specific antibody recognizing this AKT modification as a readout of PI3K-AKT pathway activation. Indeed, using immunoblotting analysis, we found that NVP-2 treatment decreased protein levels of p110α, p110β, p85α, and IRS-2 in a dose-dependent fashion. Critically, these effects were accompanied by gradual attenuation of the PI3K signaling as determined by decreasing Ser473-P AKT levels, while total levels of AKT were affected minimally. In contrast, activity of the mitogen-activated protein kinase signaling pathway which we monitored using phospho-specific and total ERK1/2 antibodies did not change, highlighting specificity of P-TEFb for the PI3K-AKT pathway. Together, we conclude that inhibition of P-TEFb hampers the oncogenic PI3K-AKT signaling cascade by repressing transcription of genes encoding key components of this pathway.

### Pharmacological targeting of the PI3K-AKT pathway elicits synthetic lethality with activators of p53

The above findings suggested that decreased activity of the pro-survival PI3K-AKT pathway is a driver of heightened dependency on P-TEFb upon activation of p53. Hence, we hypothesized that direct pharmacological targeting of the PI3K-AKT pathway elicits synthetic lethality with non-genotoxic and genotoxic activators of p53. To address this prediction, we conducted cytotoxicity assays using the parental and *TP53* null HCT116 cells which we treated with Nutlin-3a, 5-FU, and a potent and selective inhibitor of class I PI3K Pictilisib (GDC-0941) alone or in combination^56^. While all compounds provoked limited toxicity over the DMSO control when administered alone, Pictilisib treatment elicited synthetic lethality with both activators of p53 in a synergistic manner in parental but not *TP53* null cells, reaching the Bliss scores of 41 and 38 with Nutlin-3a and 5-FU, respectively (Fig. 6f and Supplementary Fig. 6a). Importantly, the lethality of non-genotoxic p53 activation and class I PI3K inhibition was abrogated in *BAX*/*BAK1* null HCT116 cells (Fig. 6f), demonstrating the dependency on the intrinsic apoptosis pathway.

To extend these findings, we next tested whether attenuating PI3K-AKT pathway triggers p53-dependent apoptosis of HCT116 cells treated with Nutlin-3a. We exposed the parental and *TP53* null HCT116 cells to increasing doses of Nutlin-3a in the absence or presence of Pictilisib and performed immunoblotting analyses to monitor apoptosis as well as activity of p53 and PI3K-AKT pathways by determining levels of cPARP, p53, p21 as well as total and Ser473-P AKT. Indeed, while Nutlin-3a treatment of the parental HCT116 cells led to stabilization of p53 and induction of p21 in dose-dependent fashion in the absence or presence of Pictilisib, cPARP levels increased only upon blocking PI3K-AKT pathway (Fig. 6g, left). We observed none of these in *TP53* null HCT116 cells (Fig. 6g, right), highlighting the requirement of p53 for this synthetic lethality. Finally, co-treatment of HCT116 with Nutlin-3a and Pictilisib led to synergistic activation of caspase 9 in BAX/BAK-dependent manner (Supplementary Fig. 6b). Together, these results demonstrate that inhibiting class I PI3K also switches the fate of Nutlin-3a-treated cells from cell-cycle arrest to apoptosis, whereby stimulation of the intrinsic apoptosis pathway plays a decisive role. Furthermore, they indicate that compromising the PI3K-AKT pathway through transcriptional repression of genes encoding key components of the pathway contributes to the synthetic lethality of PAPI.

### Synthetic lethality of p53 activation and P-TEFb inhibition remains effectual in spheroid cultures

Three-dimensional spheroid cell cultures are a widely used platform for pre-clinical assessment of anti-cancer treatments^57^. In contrast to monolayer cultures, spheroids possess several *in vivo* characteristics of tumor microenvironments, including well-organized spatial architecture of cells with different proliferation status, complex cell-cell contacts, oxygen and nutrient gradients and rewired metabolism. Therefore, we assessed whether the synthetic lethal combinations defined in this study remained effectual in HCT116 spheroids.

We established scaffold-free spheroids of about 400 μm in size and treated them for four days with DMSO, Nutlin-3a, 5-FU and NVP-2 alone and in combination as indicated (Fig. 7a). To assess the viability of spheroids, we used confocal microscopy to image spheroids stained with a cocktail of three dyes. We identified live cells with an intracellular esterase activity indicator Calcein AM emitting green fluorescence, dead cells with a cell membrane impermeable DNA stain DRAQ7 emitting red fluorescence, and total cell nuclei with the cell membrane permeable DNA stain Hoechst emitting blue fluorescence. When administered alone, all compounds had a modest impact on spheroid viability and architecture. While both 5-FU and NVP-2 treatment resulted in limited toxicity, Nutlin-3a yielded spheroids with considerably less cells, indicative of p53-dependent cell-cycle arrest. Critically, both activators of p53 sensitized the spheroids to the sub-lethal dose of NVP-2 as evidenced by massive cell death that was accompanied by the loss of spheroid architecture (Fig. 7a and Supplementary Fig. 7a). To quantify viability of spheroids, we determined a ratio between dead and live cells for each spheroid by dividing intensities of Calcein AM and DRAQ7 staining in segmented cells, which we further normalized to the total spheroid cell number determined by Hoechst staining. Corroborating the confocal images, Nutlin-3a and 5-FU synergized with NVP-2 in eliciting death of spheroids (Fig. 7b).

**Figure 7.**
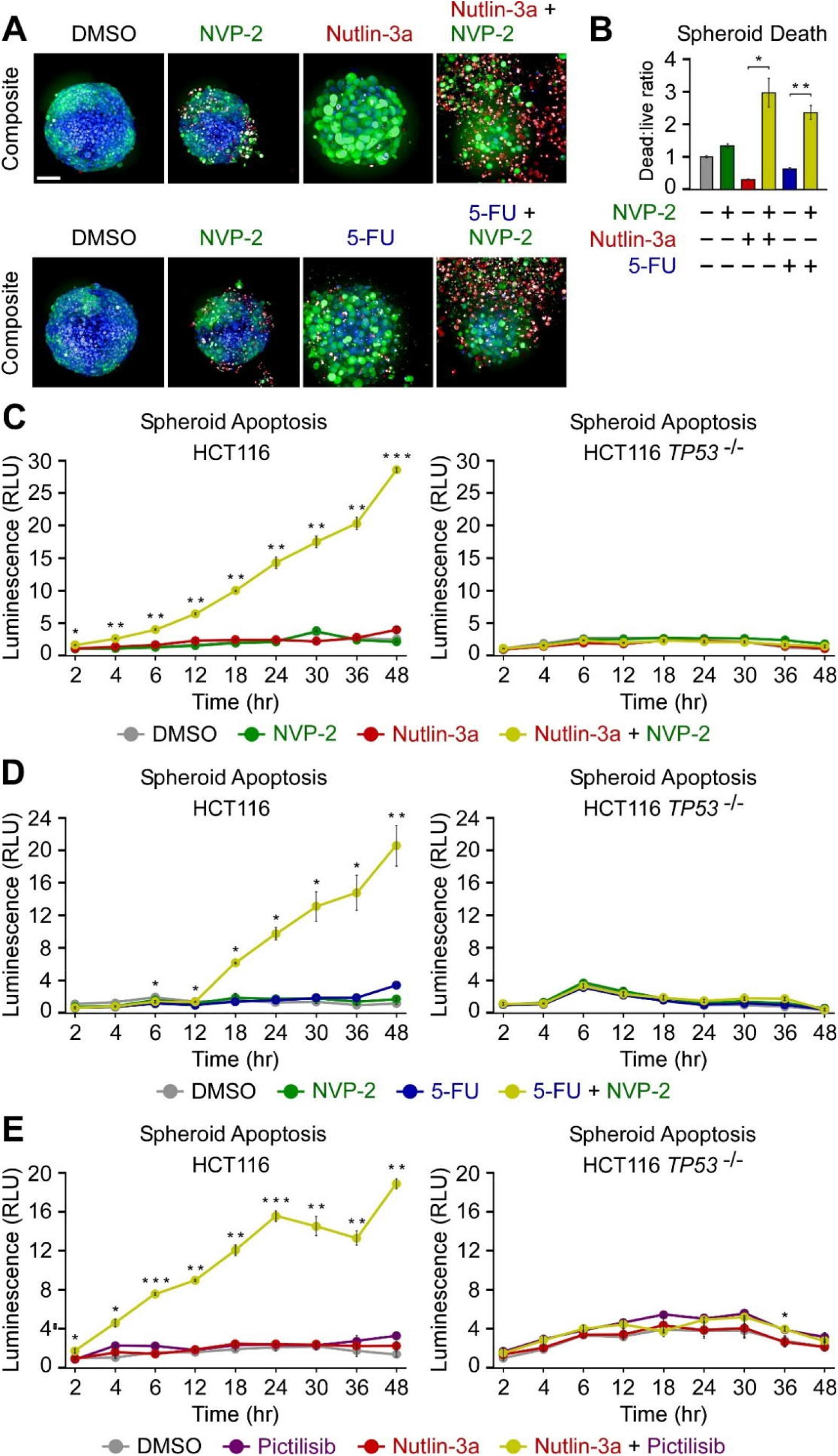
Combination treatments within the framework of p53 activation and P-TEFb inhibition trigger apoptosis of HCT116 spheroid cultures. (A) Representative composite images of HCT116 spheroid cultures treated with DMSO, NVP-2 (10 nM), Nutlin-3a (10 μM), and 5-Fluorouracil (5-FU; 25 μM) alone and in the combinations as indicated. Spheroids were formed for 48 hr, exposed to the treatments for 96 hr and stained with Calcein AM (1 μM), DRAQ7 (6 μM), and Hoechst (1x) prior to confocal microscopy imaging. Scale bar, 100 μm. (B) Quantification of death of HCT116 spheroid cultures treated with DMSO, NVP-2 (10 nM), Nutlin-3a (10 μM), and 5-Fluorouracil (5-FU; 25 μM) alone and in the combinations as indicated. Death of spheroids was determined as a ratio of integrated intensities from Calcein AM and DRAQ7 staining in segmented cells normalized to the total spheroid cell number quantified by Hoechst staining (dead:live ratio). The results presented as dead:live ratio values relative to the values of DMSO-treated cells are plotted as the mean ± s.e.m. (n = 3). *, P < 0.05; **, P < 0.01, determined by Student’s *t* test using Nutlin-3a and Nutlin-3a + NVP-2, and 5-FU and 5-FU + NVP-2 data sets. (C-E) Apoptosis of HCT116 and HCT116 *TP53* ^-/-^ cell spheroid cultures treated with DMSO (grey), NVP-2 (10 nM; green), Nutlin-3a (10 μM; red), 5-Fluorouracil (5-FU; 25 μM) (blue) and Pictilisib (0.5 μM; magenta) alone and in combination (gold). Spheroids were formed for 48 hr prior to the treatments. Results obtained at the time points indicated below the graphs using RealTime-Glo Annexin V Apoptosis and Necrosis Assay are presented as luminescence values relative to the values of DMSO-treated cells at 2 hr and plotted as the mean ± s.e.m. (n = 3). *, P < 0.05; **, P < 0.01; ***, P < 0.001, determined by Student’s *t* test using Nutlin-3a and Nutlin-3a + NVP-2 (C), 5-FU and 5-FU + NVP-2 (D), and Nutlin-3a and Nutlin-3a + Pictilisib (E) data sets.

Finally, we conducted quantitative kinetic measurements to monitor apoptosis of spheroids exposed to combination treatments defined in this study as synthetic lethal using monolayer cultures. We detected negligible levels of apoptosis when analyzing HCT116 spheroids treated for three days with Nutlin-3a, 5-FU, NVP-2, and Pictilisib. In contrast, the respective combination treatments elicited robust apoptosis in a highly synergistic manner (Fig. 7c-e, left). As anticipated, p53 was required for this effect as the combinations failed to result in apoptosis of *TP53* null HCT116 spheroids (Fig. 7c-e, right). Likewise, Nutlin-3a treatment sensitized HCT116 *TP53* ^R248/+^ but not p53-deficient HCT116 *TP53* ^R248/-^ spheroids to NVP-2 (Supplementary Fig. 7b). In addition, both 5-FUR and 5-FdUR, the ribose and deoxyribose derivatives of 5-FU, elicited apoptosis in the presence of NVP-2 (Supplementary Fig. 7c, d). Finally, within the canonical PAPI combination treatment, we replaced P-TEFb inhibitor NVP-2 with DT-061, a prototypical small-molecule activator of PP2A that stabilizes and allosterically activates a specific heterotrimeric PP2A holoenzyme which dephosphorylates key P-TEFb substrates including SPT5 and the CTD of Pol II^9,58^. We postulated that PP2A activation should akin to P-TEFb inhibition synergize with p53 activation in inducing apoptotic cell death. Indeed, non-genotoxic and genotoxic activation of p53 by Nutlin-3a and 5-FU, respectively, sensitized HCT116 spheroids to apoptosis by PP2A activator DT-061 (Supplementary Fig. 7e, f), illustrating possibilities to expand further the arsenal of synthetic lethal combinations of PAPI. Together, we conclude that the synthetic lethal combination treatments uncovered with monolayer cell cultures remained effectual in a translational spheroid setting.

## DISCUSSION

The central role of tCDKs in gene control and their emergence as vital players in increasing number of cancer types has elevated the interest in these kinases as potential therapeutic targets. In this study, we harnessed a comprehensive oncology library to identify compounds that rendered colon cancer-derived cells dependent on Pol II transcription driven by P-TEFb. Amongst the screen hits, we focused on antimetabolites and MDM2 inhibitors, the broadly used chemotherapeutics and small-molecule PPI inhibitors with a huge clinical potential, respectively, and uncovered a set of small molecule combination treatments that triggered cell death by apoptosis in synergistic and p53-dependent fashion. Importantly, we revealed that stimulation of the intrinsic apoptosis pathway drives cellular hypersensitivity to P-TEFb inhibition upon non-genotoxic stabilization of p53. Finally, we showed that combination treatments dismantled cancer cell spheroids by apoptosis, offering a rationale for further studies to assess effectiveness of the combinations in translational settings.

Our study contributes to the growing repertoire of synthetic lethal interactions with inhibitors of P-TEFb. Previous work identified genetic and epigenetic alterations in cancer cells that heighten the reliance on P-TEFb. The prime examples are amplification of *MYC* and rearrangement of *MLL1*, whereby both oncoproteins redirect P-TEFb to their loci to drive oncogene-specific transcription programs, providing an explanation for increased sensitivity to P-TEFb inhibitors^17^. In addition, perturbation of several targets, including BRD4^35,59^, PARP^60^, and CDK2^61^ was reported to elicit lethality of cancer cells in combination with P-TEFb inhibitors. However, many of these studies employed inhibitors exhibiting sub-optimal selectivity towards P-TEFb, entailing a caveat that targeting of additional kinases contributed to the lethality. By conducting our screen with the highly selective inhibitor NVP-2, we here validated some of the previous reports and identified novel synthetic lethal interactions. We revealed further that inhibiting class I PI3K and activating PP2A also sensitizes cancer cells to the representative p53-activating compounds Nutlin-3a and 5-FU. As PI3K and PP2A are important targets in cancer^52,62^, our study offers new opportunities for the rational design of combination therapies that rely on functional p53.

Mechanistically, we uncovered a molecular basis that underpins the ability of P-TEFb inhibitor NVP-2 to switch the fate of Nutlin-3a-treated cells from cell-cycle arrest to apoptosis. Our evidence supports a model that differential sensitivity of two key sets of genes to P-TEFb inhibition enables the fate switching. On the one hand, mRNA levels of p53-induced pro-death genes of the intrinsic apoptosis pathway remain largely unhindered in the face of sub-lethal inhibition of P-TEFb, which is critical to the onset of apoptosis. As illustrated by our transcription shutdown experiments using triptolide or high concentration of NVP-2, cells require active Pol II transcription to execute apoptosis. On the other hand, however, mRNA levels of pro-survival genes that code for key components of the oncogenic PI3K-AKT signaling pathway decreased under the inhibition of P-TEFb. Of note, PI3K-AKT pathway antagonizes apoptosis by phosphorylation and inhibition of pro-apoptotic mediators BAD and caspase 9, wherein AKT acts to release pro-survival BCL-2/BCL-X_L_ from BAD and to prevent cytochrome *c*-mediated activation of caspase 9, respectively^55^. Therefore, we propose that following p53 activation, P-TEFb suppresses intrinsic apoptosis pathway in part by facilitating the signaling through the PI3K-AKT cascade at the level of the class IA PI3K gene transcription. Given the centrality of P-TEFb in transcription by Pol II ^5^, it is likely that its targeting impacts additional genes of which altered expression contributes to the activation of apoptosis.

Our work provides grounds for improving antimetabolite-based chemotherapeutic interventions with the addition of highly selective antagonists of P-TEFb. Importantly, it promises to reinvigorate the ongoing efforts of reactivating tumor suppressive capabilities of p53 with Nutlin-3a-based combination therapies^29^. Since the discovery of Nutlin-3a nearly twenty years ago, there has been a tremendous interest in delivering this class of non-genotoxic activators of p53 into the clinic. While up to a half of cancers carry *TP53* loss-of-function mutations in both alleles, the remaining half keep p53 in check through other means such as *MDM2* amplification, making WT p53 an appealing target for therapeutic intervention. However, Nutlin-3a and its derivatives elicit reversible cell-cycle arrest but not apoptosis in most cancer cell systems examined^32^. Therefore, our findings on the fate switching of Nutlin-3a-treated cells by inhibitors of P-TEFb could come to the rescue. Because small-molecule MDM2 degraders are presumed to exhibit increased potency and therapeutic index over MDM2 inhibitors^63^, combining the degraders with highly selective P-TEFb and PI3K antagonist or PP2A agonists warrants further investigation.

For over three decades, a goal of clinical oncology has been the development of therapies that promote elimination of cancer cells by apoptosis^24^. Interestingly, our screen also identified SMAC mimetics amongst the top compounds that cooperate with NVP-2 in cancer cell killing. By mimicking the action of SMAC that gets released into the cytosol during MOMP, SMAC mimetics target pro-survival IAP proteins to facilitate cell death by promoting caspase 3, 7, and 9 activity^64^. We therefore envision that targeting P-TEFb in combination with many suppressors of intrinsic apoptosis pathway could become a viable therapeutic strategy for irreversible demise of cancer cells.

## METHODS

### Key Resources

A list of key resources is provided in Supplementary Table 1a-e.

### Cell Culture

HCT116 parental cell line and its derivatives were cultured in McCoy’s 5A Medium (Sigma) and Dulbecco‘s Modified Eagle‘s Medium (Sigma) supplemented with 10% FBS and 100 U/ml penicillin/streptomycin. CCD 841 CoN cell line was cultured in Eagle’s Minimum Essential Medium (Lonza) supplemented with 10% FBS and 100 U/ml penicillin/streptomycin. All cell lines were maintained at 37°C with 5% CO2. Cell lines were confirmed regularly to be mycoplasma-free using a qPCR-based Mycoplasmacheck detection kit (Eurofins). Cell lines were not authenticated by us, but retrieved from trusted sources as listed in Supplementary Table 1d.

### Chemicals

NVP-2, Nutlin-3a, 5-Fluorouridine, 5-Fluorodeoxyuridine, THZ531, and Pictilisib were from MedChemExpress. 5-fluorouracil, Senexin A, OTS964, and Triptolide were from Selleck Chemicals. YLK-5-124 and THAL-SNS-032 were a gift from Nathanael S. Gray’s Laboratory (Stanford University, USA). iCDK9 was a gift from Qiang Zhou’s Laboratory (University of California, Berkley, USA). PP2A activator DT-061 was a gift from Jukka Westermarck’s Laboratory (University of Turku, Finland).

### FO5A Oncology Library Screen and Data Analysis

FO5A version of the FIMM oncology collection was obtained from various commercial chemical vendors^65^. The library comprised 527 single compounds and 1 drug combination, of which 28% represented approved anticancer drugs, 55% - emerging investigational compounds, and 17% - chemical probes covering a wide range of molecular targets^65^ (Supplementary Fig. 1a). Drug Sensitivity and Resistance Testing was performed as described previously^65^ with minor modifications. Cytotoxicity was evaluated using CellTox Green Cytotoxicity Assay (Promega), whereby CellTox Green Dye was added to suspension of DMSO- and NVP-2-treated HCT116 cells that were seeded at 2,000 cells/well on the 384-well plates containing the library’s compounds in 25 μl final volume using a BioTek MultiFlo FX microplate dispenser. After 72 hr of incubation, fluorescence was measured with a Pherastar FS multimode plate reader (BMG Labtech) according to manufacturer’s instructions. The cell viability data were processed using an in-house developed drug screening data analysis application Breeze^66^. The data were normalized to negative control wells containing DMSO and positive control wells containing 100 μmol/L benzethonium chloride that kills all cells. Drug sensitivity scores (DSS)^67^ were calculated for each treatment. Differential DSS (dDSS) was further calculated by subtracting DSS values of NVP-2-containing plate sets from the values of DMSO-containing plate sets. Positive dDSS values indicated that NVP-2 enhances the cytotoxic effect of a given compound, while negative dDSS values suggest the opposite effect. The waterfall plots showing compounds having significant differences in dDSS across the screens were generated using R programming language with the help of package ggplot2.

### Cytotoxicity, Viability, Apoptosis, and Caspase 9 Activity Assays

Cytotoxicity, viability, apoptosis, and Caspase 9 activity assays were conducted using CellTox Green Cytotoxicity Assay (Promega), CellTiter-Glo 2.0 Cell Viability Assay (Promega), RealTime-Glo Annexin V Apoptosis and Necrosis Assay, and Caspase-Glo 9 Assay, respectively, according to manufacturer’s instructions. For cytotoxicity, viability, and apoptosis assays, HCT116 parental cell line and its derivatives were seeded at 20,000 cells/well on 96-well plates for 16 hr and treated with the indicated compound combinations and corresponding volume of DMSO. For Caspase 9 activity assay, the parental and *BAX*/*BAK* ^−/−^ HCT116 cells were seeded at 8,000 cells/well on 384-well plates for 6 hr and treated with the indicated compound combinations. Fluorescence and luminescence measurements were taken at the indicated time points using PerkinElmer Victor X3 reader. Results from at least three independent experiments are presented as values relative to the values of DMSO-treated cells and plotted as the mean ± SEM. For determining IC_50_ value of NVP-2 in the presence and absence of Nutlin-3a and 5-FU, HCT116 cells were seeded at 5,000 cells/well on 96-well plates for 6 hr and treated with the indicated compounds for 72 hr, after which cell viability was determined using CellTiter-Glo 2.0 Cell Viability Assay (Promega). Luminescence measurements were taken using PerkinElmer Victor X3 reader, and results from two independent experiments were normalized to the values of DMSO-treated cells and plotted as the average ± s.d.

### Data Analysis of Cytotoxicity and Viability Matrix Assays

To obtain % of inhibition in cytotoxicity and viability assays, the raw fluorescence and luminescence values are presented as a ratio to the DMSO-treated control, normalized to the difference between DMSO and positive control sample representing maximum cytotoxicity. Maximum cytotoxicity was defined as a maximum fluorescence value obtained from the wells showing absolute cell death. An average of three independent experiments was plotted using the Seaborn Python data visualization library (https://github.com/mwaskom/seaborn). Bliss synergy scores were calculated using Synergyfinder^34^.

### Western Blotting Assay

Whole-cell extracts (WCE) were prepared using lysis buffer C (20 mM Tris-HCl, 0.5% NP-40, 150 mM NaCl, 1.5 mM MgCl2, 10 mM KCl, 10% Glycerol, 0.5 mM EDTA, pH 7.9) on ice for 1 hr in the presence of EDTA-free Protease Inhibitor Cocktail (Sigma). Lysates were cleared by centrifugation at 20,000 g for 15 min, boiled in SDS running buffer supplied with 10% of β-Mercaptoethanol for 5 min, separated using 10% SDS-PAGE, and transferred to nitrocellulose membrane. For immunoblotting, the following antibodies were used according to manufacturers’ instructions: anti-CDK9 (Santa Cruz Biotechnology, 1:4000); anti-p53 (Santa Cruz Biotechnology, 1:3000); anti-Cleaved PARP (Cell Signaling Technology, 1:3000); anti-p21 (Santa Cruz Biotechnology, 1:4000); anti-GAPDH (Santa Cruz Biotechnology, 1:10,000); anti-RNA polymerase II CTD repeat YSPTSPS (phospho S2) (Abcam, 1:3000); anti-RNA polymerase II RPB1 (Santa Cruz Biotechnology, 1:1000); anti-p110α (Santa Cruz Biotechnology, 1:100); anti-p110β (Santa Cruz Biotechnology, 1:500); anti-p85α (Santa Cruz Biotechnology, 1:3000); Anti-IRS-2 (Santa Cruz Biotechnology, 1:1000); anti-phospho-Akt (Ser473) (Cell Signaling Technology, 1:4000); anti-pan-Akt (Santa Cruz Biotechnology, 1:4000); anti-phospho-p44/42 MAPK (Erk1/2) (Thr202/Tyr204) (Cell Signaling Technology, 1:4000); anti-p44/42 MAPK (Erk1/2) (Cell Signaling Technology, 1:4000). Manufacturers provide validation for all antibodies.

### RNA Sequencing

Total RNA isolation and on-column digestion of genomic DNA was performed using the RNeasy Mini Kit (Qiagen) according to the manufacturer’s instructions. RNA quality was assessed by TapeStation 4200 and samples with a RIN value of 10 were used for library preparation and sequencing. PolyA-selected libraries were made using NEBNext Poly(A) mRNA Magnetic Isolation Module and NEBNext Ultra Directional RNA Library Prep Kit (New England Biolabs) according to the manufacturer’s instructions. The libraries were pooled and sequenced on one lane of an Illumina NextSeq500 (High Output 1 × 75 bp; single-end).

### Analysis of RNA Sequencing Datasets

RNA-seq reads were mapped against the human genome (hg38) and human rRNA sequences with ContextMap version 2.7.9^68^ using Burrows-Wheeler Alignment tool^69^. Number of read counts per gene and exon were determined from the mapped RNA-seq reads in a strand-specific manner using featureCounts^70^ and gene annotations from Gencode version 25^71^. Differential gene expression analysis was performed using edgeR^72^. P-values were adjusted for multiple testing using the method by Benjamini and Hochberg^73^ and genes with an adjusted p-value ≤ 0.001 were considered differentially expressed. For further analysis, we included only genes expressed (Reads per kilo base per million mapped reads (RPKM) >=1) in at least condition. Analysis workflows were implemented and run using the Watchdog workflow management system^74^. Volcano plots and heatmaps were created in R (https://www.R-project.org/). Hierarchical clustering analysis of genes in heatmaps was performed using Ward’s clustering criterion^75^ and Euclidian distances. Gene set enrichment analysis on log2 fold-changes for protein-coding genes was performed using the command line tool from http://www.gsea-msigdb.org/ (version 4.0.2)^44^ and hallmark gene sets from Molecular Signatures Database.

### RNA Extraction and RT-qPCR Assay

HCT116 cells were seeded on 6-well plates for 16 hr and treated at 80% confluency with the indicated compound combinations and corresponding volume of DMSO. RNA samples were extracted using TRI Reagent (Sigma), DNase-treated with the Turbo DNA-Free kit (Thermo Fisher Scientific), and reverse transcribed with M-MLV reverse transcriptase (Thermo Fisher Scientific) and random hexamers (Thermo Fisher Scientific) according to the manufacturers’ instructions. qPCR reactions were performed with diluted cDNAs, primer pairs that spanned exon-exon junctions, and FastStart Universal SYBR Green QPCR Master (Rox) (Sigma) using Roche LightCycler 480. Primers were from Integrated DNA Technologies and designed using PrimerQuest Tool. Results from three independent experiments were normalized to the values of *GAPDH* mRNA and DMSO-treated cells, and plotted as the mean ± s.e.m. Sequences of the primers used are listed in Supplementary Table 2.

### Quantitative Chromatin Immunoprecipitation Assay

ChIP-qPCR assay was performed as described previously^76^ with the following modifications. HCT116 cells cultured on 15 cm plates were treated at approximately 80% confluency with DMSO or 20 nM NVP-2 for 3 hr. Sixteen million cells were cross-linked with formaldehyde and the cell pellets were lysed in 800 μL of RIPA buffer (50 mM Tris pH 8.0, 150 mM NaCL, 5 mM EDTA pH 8.0, 0.5% sodium deoxycholate, 1% NP-40, 0.1% SDS) in the presence of EDTA-free Protease Inhibitor Cocktail (Sigma). Lysates were sonicated with 12 cycles of 14 s sonication with the Misonix XL-2000 Ultrasonic Liquid Processor using the P-1 Microprobe 3.2 mm tip, power setting 10, in 1.7 mL

RNase/DNase-free microcentrifuge tubes (Sigma), which were kept for 1 min on ice between the cycles. After centrifugation at 13,000 g for 15 min, 30 μL of the cleared chromatin was stored at −80°C for determining DNA input. The rest of the sample was divided in four equal parts, which were supplemented with additional 800 μL of RIPA buffer and incubated overnight at 4°C with 15 μL of antibody-coupled protein G Dynabeads (Thermo Fisher Scientific). Before adding the cleared chromatin, the beads were pre-blocked with bovine serum albumin and salmon sperm DNA overnight at a final concentration of 0.2 μg/μl, pre-incubated in 500 μL of RIPA buffer for 4 h with the antibody and collected by magnetic stand to remove the unbound antibody. We used 2 μg of anti-RNA polymerase II RPB1 NTD (Cell Signaling Technology), 1 μg of anti-RNA polymerase II CTD repeat YSPTSPS (phospho S2) (Abcam), and 1 μg of anti-RNA polymerase II CTD repeat YSPTSPS (phospho S5) (Abcam) antibodies. Additionally, 2 μg of normal rabbit IgG (Santa Cruz Biotechnology) antibodies were used to determine specificity of the signals. Validation for all antibodies is provided on the manufacturers’ websites. ChIP samples were dissolved in 30 μL of water and 0.4 μL of the solution was used per one qPCR reaction. DNA input was dissolved in 100 μL of water, diluted 50 times, and 1 μL of the solution was used per one qPCR reaction. Samples were amplified using FastStart Universal SYBR Green QPCR Master (Rox) (Sigma), DNA-specific primer pair, and LightCycler 480 II (Roche) machine. Primers were from Integrated DNA Technologies and designed using PrimerQuest Tool. Values were normalized to their levels in DNA inputs and the values of DMSO-treated cells. Ser2-P and Ser5-P ChIP values were further normalized to the values of total Pol II. Results from two independent experiments are presented as the mean ± s.e.m. Sequences of the primers used are listed in Supplementary Table 3.

### Confocal Imaging of HCT116 Spheroids

HCT116 cell were seeded on 384-well, ultra-low attachment, Corning U-bottom, black clear-bottom imaging plates (Corning) at density of 400 cells/well and spun down at 200g for 10 minutes. After culturing the cells for 48 hr, the spheroids were treated with the indicated compound combinations for 96 hr. Spheroids were stained with a freshly prepared PBS solution of 1 μM Calcein AM, 6 μM of DRAQ7, and 1x Hoechst 33342 (Invitrogen) for 6 hr, after which imaging was performed with Opera Phenix confocal spinning-disk high-content screening microscope (PerkinElmer). Screening was conducted with a 40x water immersion objective. Fields of view were automatically selected from a pre-scan image with a 10% overlap using 90 pre-determined Z focus planes with laser-based autofocusing. The images were captured with two Andor Zyla sCMOS cameras (Andor Technology). 3D Image analysis was performed with Harmony v4.9 High-Content Imaging and Analysis Software (PerkinElmer). Cells were segmented according to the Hoechst 33342 nuclear staining with a cytoplasmic mask from combined channel images. Results were quantified as a dead:live ratio of integrated intensities from DRAQ7 and Calcein AM staining in segmented cells and normalized against the observed cell number as determined by Hoechst 33342 staining. Results of the normalized dead:live ratio values from three independent experiments were further normalized to the values of DMSO-treated cells and plotted as the mean ± s.e.m.

### Quantification and Statistical Analysis

Differential gene expression analysis of RNA-seq data was performed using edgeR^72^. P-values obtained from edgeR were corrected by multiple testing using the method by Benjamini and Hochberg^73^ for adjusting the false discovery rate (FDR) and a p-value cutoff of 0.01 was applied. Data shown for all qPCR-based experiments and functional assays were collected from at least 3 biological replicates as indicated in individual figure legends and are presented as means ± s.e.m. Statistical significance and *P* values were determined by Student’s *t* test performed between the indicated paired groups of biological replicates. Where the results were not statistically significant, *P* values are not indicated.

## ACKNOWLEDGEMENTS

We thank Joaquin M. Espinosa for sharing parental and HCT116 *TP53* ^-/-^ cell lines; Bert Vogelstein for sharing HCT116 *TP53* ^R248W/+^ and HCT116 *TP53* ^R248W/+^ cell lines; Ana J. Garcia-Saez for sharing HCT116 *BAX*/*BAK1* ^-/-^ cell line; Qiang Zhou for sharing i-CDK9; Nathanael S. Gray for sharing YLK-5-124 and SNS032-THAL; Jukka Westermarck for sharing DT-061; FIMM High Throughput Biomedicine Unit for executing the FO5A screen and data analysis; FIMM High Content Imaging and Analysis Unit for confocal imaging and analysis; Biomedicum Functional Genomics Unit at the University of Helsinki for RNA-seq. This work was funded by Sigrid Juselius Foundation [4707979 to M.B.]; Academy of Finland [1309846 to M.B.]; Cancer Society of Finland [4708690 to M.B.]; Deutsche Forschungsgemeinschaft (DFG) [FR2938/9-1 and FR2938/10-1 to C.C.F.].

## AUTHOR CONTRIBUTIONS

M.B. and A.B. conceptualized the study and were assisted by Z.W and M.M. in experimental design. M.B. supervised the study. S.G.K. designed and executed the FO5A screen with the assistance of M.B. S.P. analyzed the FO5A screen data and generated the associated waterfall plots. A.B. performed and analyzed all combinatorial titration cytotoxicity and viability assays and generated the associated heatmaps. M.M. conducted caspase 9 activity and cytotoxicity assays in the parental and HCT116 *BAX*/*BAK1* ^-/-^ cell lines, and spheroid viability experiments. A.H. imaged the spheroids and analyzed the data. T.L. performed ChIP-qPCR assays. C.C.F. analyzed RNA-seq data and generated the associated volcano plots and heatmaps. All other experiments were performed and analyzed by Z.W. M.B. wrote the manuscript.

## Supplementary Information

**Supplementary Figure 1.**
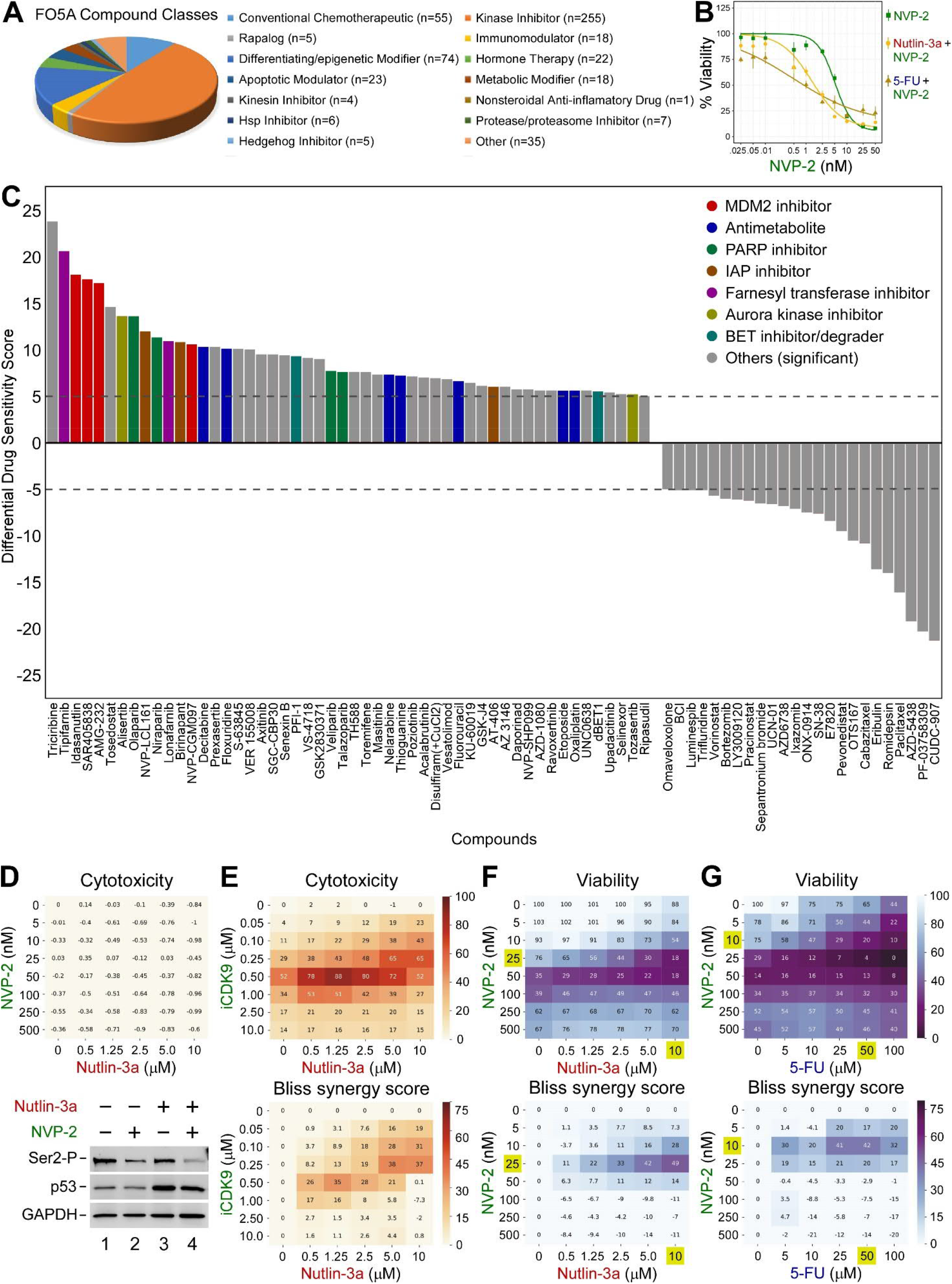
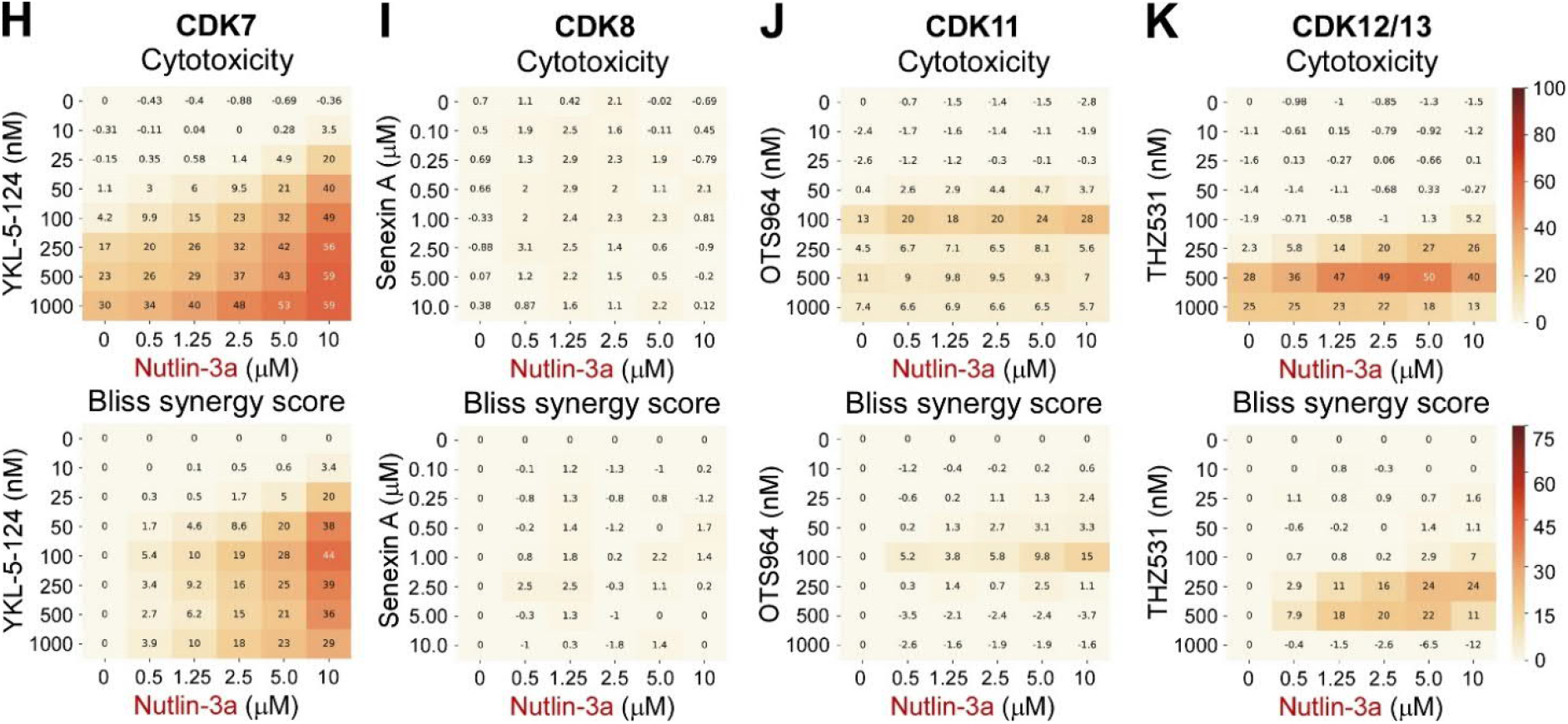
Selective inhibition of CDK7 and P-TEFb but not other tCDKs is synthetic-lethal with MDM2 inhibitor Nutlin-3a. (A) Compound classes of the FO5A oncology library. Pie chart represents classes of investigational and clinical anti-cancer compounds listed on the right. (B) Dose-response viability curves of HCT116 cells treated with increasing doses of NVP-2 (green) alone and in combination with Nutlin-3a (10 μM; light gold) or 5-Fluorouracil (5-FU; 25 μM, dark gold) as indicated. Viability values obtained at 72 hr of the treatments using CellTiter-Glo 2.0 Cell Viability Assay were normalized to the values of DMSO-treated cells and are presented as percentages of the maximum viability which was set at 100 %. Results are presented as the average ± s.d. (n = 2). (C) Waterfall plot representing results of the screen. Compounds with dDSS values ≥ 5 and ≤ -5 were considered as significant and depicted according to the legend on the right. (D) (Top) 8 × 6 cytotoxicity matrix with combinatorial titrations of NVP-2 (green) with Nutlin-3a (red) at indicated doses in non-transformed CCD 841 CoN colon epithelial cells. Cytotoxicity values obtained at 48 hr of the treatments using CellTox Green Cytotoxicity Assay were normalized to the values of DMSO-treated cells and are presented as percentages of the maximum cytotoxicity which was set at 100 %. Results represent the average of independent experiments (n = 3). (Bottom) CCD 841 CoN cells were treated with the indicated combinations of NVP-2 (10 nM) and Nutlin-3a (10 μM) for 8 hr prior to preparation of whole cell extracts and detection of the indicated proteins by Western blotting. (E, H-K) 8 × 6 matrices with combinatorial titrations of iCDK9 (green) and the indicated tCDK inhibitors (black) with Nutlin-3a (red) at indicated doses to test for the synthetic lethality of compounds in HCT116 cells, depicting cytotoxicity (top) and synergy (bottom) of the combinations. Targeted tCDKs are indicated on top. Cytotoxicity values obtained at 48 hr of the treatments using CellTox Green Cytotoxicity Assay were normalized to the values of DMSO-treated cells and are presented as percentages of the maximum cytotoxicity which was set at 100 %. Results represent the average of independent experiments (n = 3). (F,G) 8 × 6 matrices with combinatorial titrations of NVP-2 (green) with Nutlin-3a (red) and 5-Fluorouracil (5-FU; blue) at indicated doses to test for the synthetic lethality of compounds in HCT116 cells, depicting viability (top) and synergy (bottom) of the combinations. Viability values obtained at 48 hr of the treatments using CellTiter-Glo 2.0 Cell Viability Assay were normalized to the values of DMSO-treated cells and are presented as percentages of the maximum viability which was set at 100 %. Results represent the average of independent experiments (n = 3). Combinations with the highest Bliss synergy scores in HCT116 cells are highlighted (gold).

**Supplementary Figure 2.**
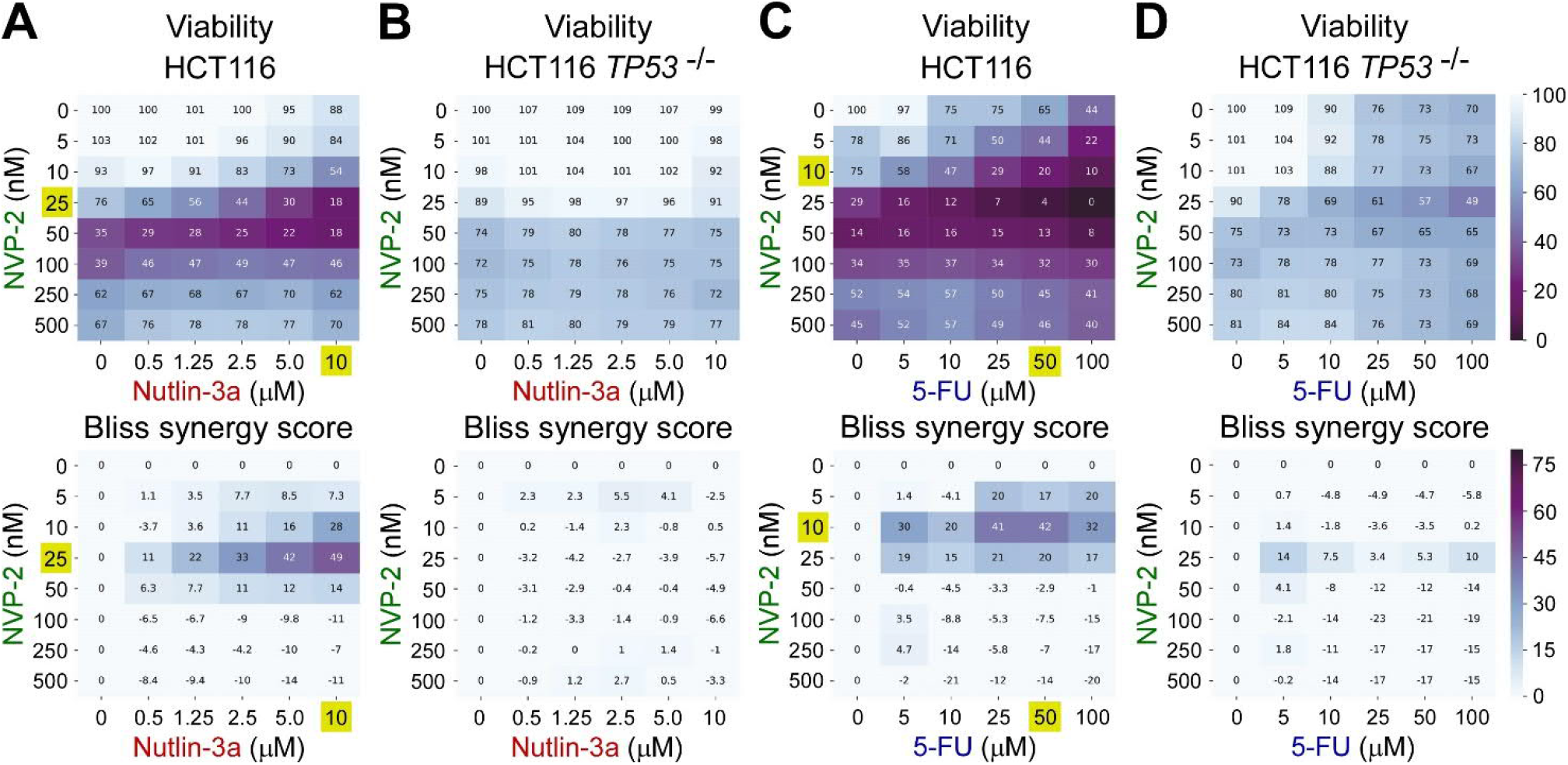
Synthetic lethality of Nutlin-3a and antimetabolites with P-TEFb inhibitor NVP-2 is p53-dependent. (A-D) 8 × 6 matrices with combinatorial titrations of NVP-2 (green) with Nutlin-3a (red) and 5-Fluorouracil (5-FU; blue) at indicated doses to test for the synthetic lethality of compounds in HCT116 and HCT116 *TP53* ^-/-^ cells, depicting viability (top) and synergy (bottom) of the combinations. Viability values obtained at 48 hr of the treatments using CellTiter-Glo 2.0 Cell Viability Assay were normalized to the values of DMSO-treated cells and are presented as percentages of the maximum viability which was set at 100 %. Results represent the average of independent experiments (n = 3). Combinations with the highest Bliss synergy scores in HCT116 cells are highlighted (gold).

**Supplementary Figure 3.**
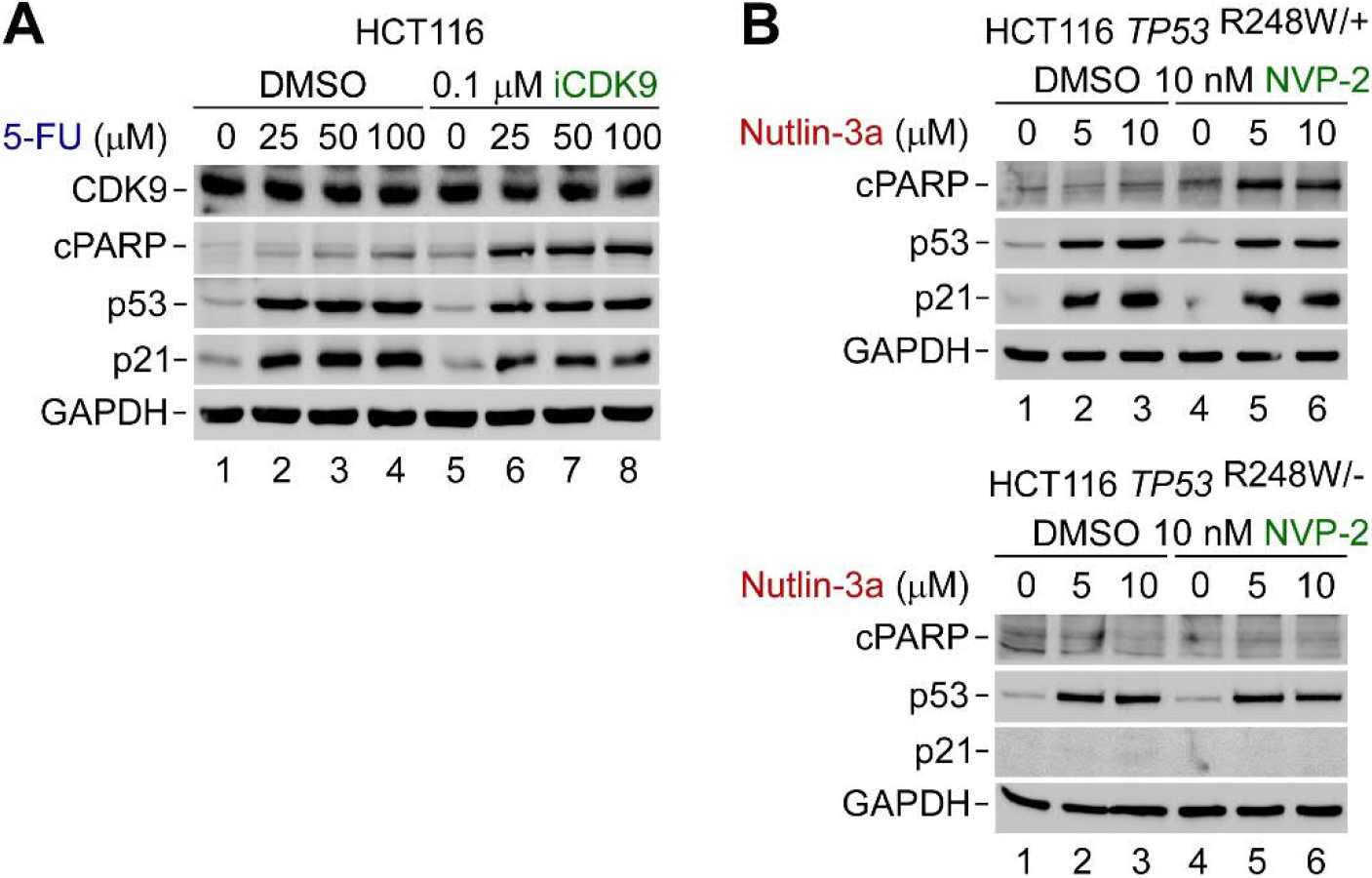
Activation of apoptosis underlies p53-dependent synthetic lethality of p53 activation and P-TEFb inhibition. (A,B) HCT116, HCT116 *TP53* ^R248W/+^ and HCT116 *TP53* ^R248W/-^ cells were treated with DMSO and indicated combinations and doses of CDK9 inhibitors (green), Nutlin-3a (red), and 5-Fluorouracil (5-FU; blue) for 24 hr prior to preparation of whole cell extracts and detection of the indicated proteins by Western blotting.

**Supplementary Figure 4.**
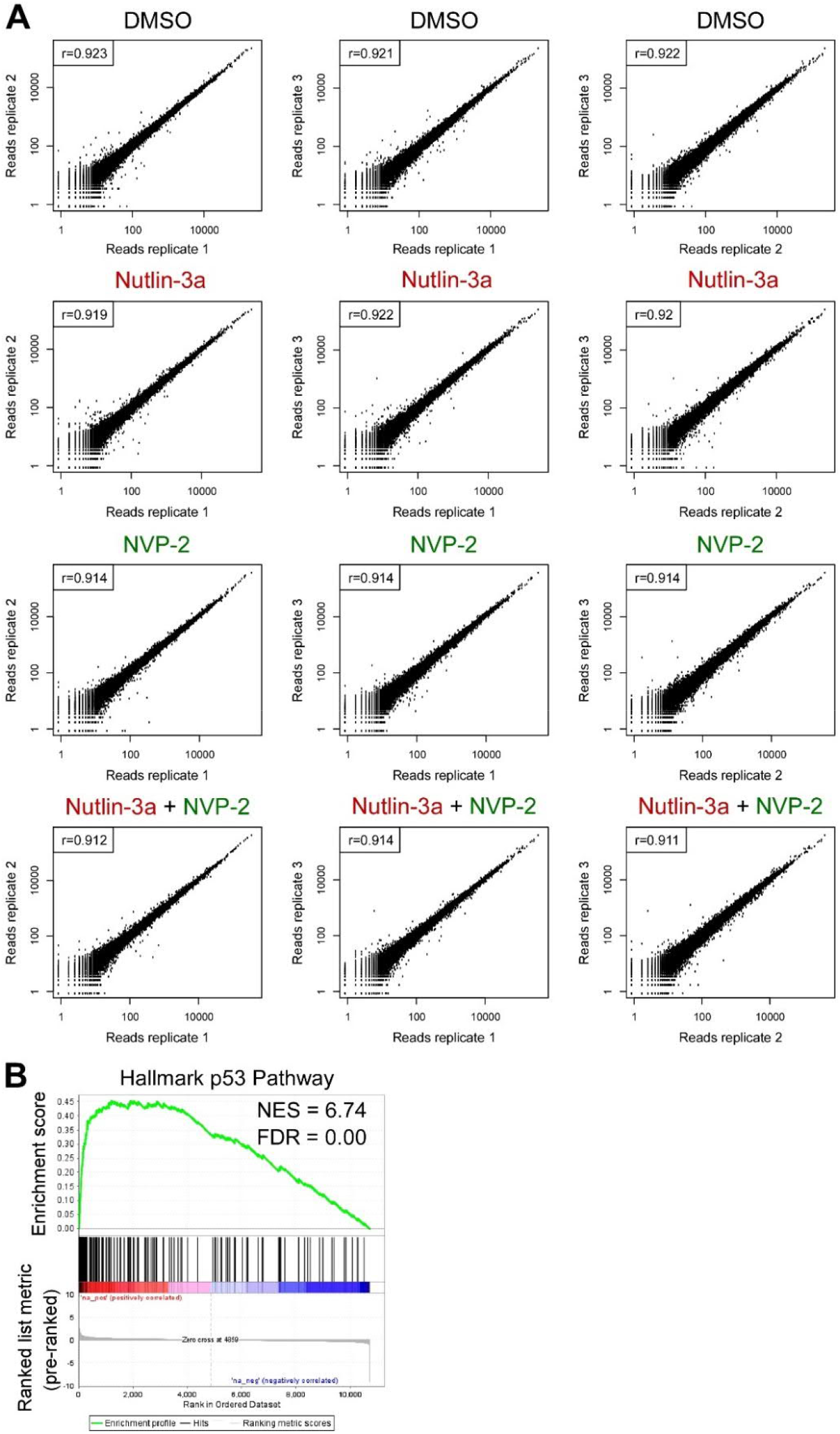
Reproducibility of RNA-seq replicates and top gene set elicited by Nutlin-3a. (A) Scatterplots comparing raw read counts of all annotated genes from RNA-seq data sets (n = 3) are shown and spearman correlation (r) is indicated. HCT116 cells were treated for eight hours as indicated on top of each graph. (B) Gene set enrichment analysis of the protein-coding gene data set regulated by Nutlin-3a. The top gene set is shown. NES, normalized enrichment score. FDR, false discovery rate.

**Supplementary Figure 5.**
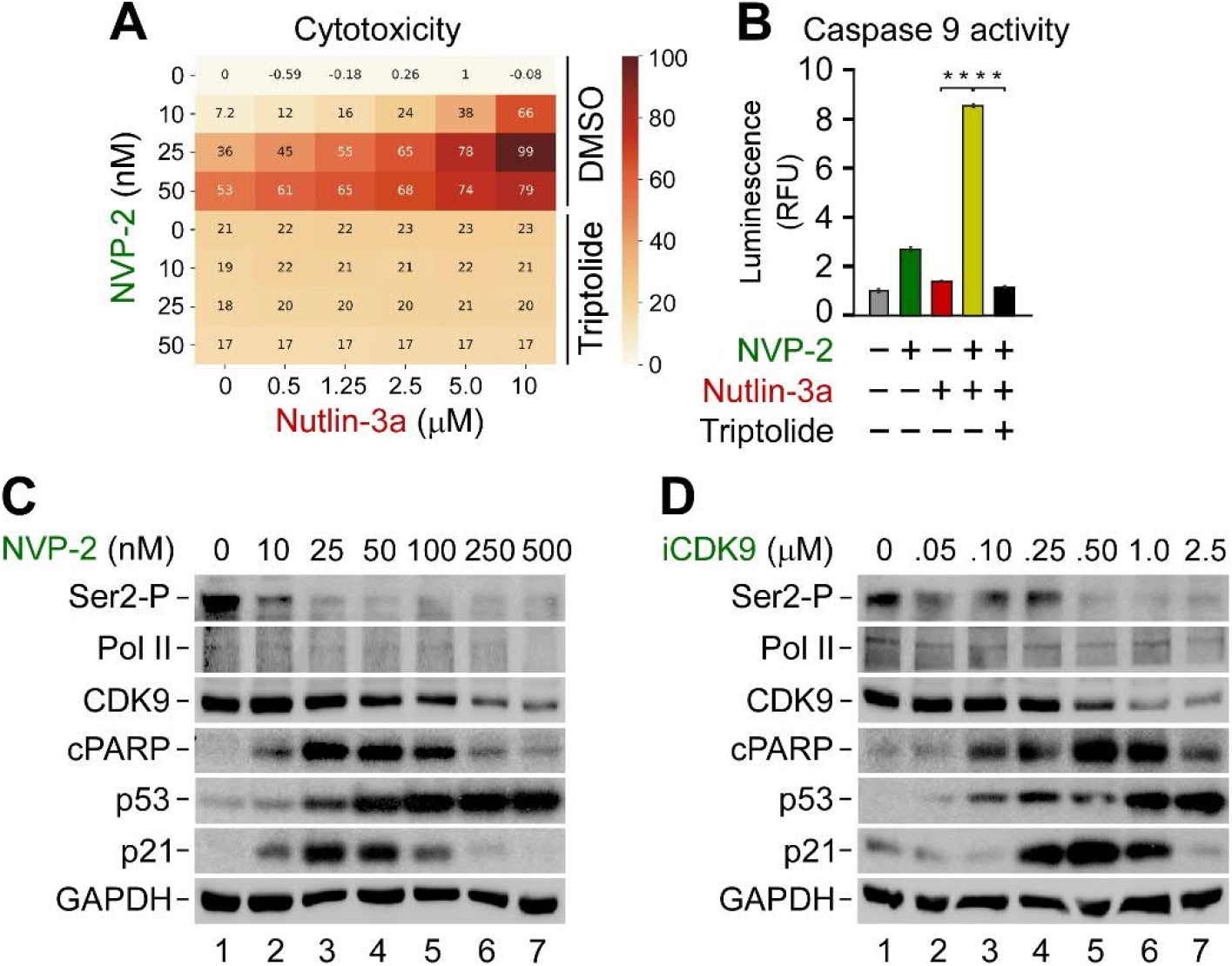
Synthetic lethality of non-genotoxic p53 activation and P-TEFb inhibition depends on intrinsic apoptosis pathway and on-going Pol II transcription. (A) 4 × 6 cytotoxicity matrices with combinatorial titrations of Nutlin-3a (red) with NVP-2 (green) at indicated doses of HCT116 cells co-treated with DMSO or Triptolide (1 μM) as indicated. Cytotoxicity values obtained at 48 hr of the treatments using CellTox Green Cytotoxicity Assay were normalized to the values of DMSO-treated cells and are presented as percentages of the maximum cytotoxicity which was set at 100 %. (B) Activity of Caspase 9 measured using Caspase-Glo 9 Assay in whole cell extracts of HCT116 cells treated with DMSO (grey), NVP-2 (3 nM; green) and Nutlin-3a (10 μM; red) alone and in combination (gold), and with Triptolide (1 μM; black) as indicated for 18 hr. Results are presented as luminescence values relative to the values of DMSO-treated cells and plotted as the mean ± s.e.m. (n = 4). **, P < 0.01, determined by Student’s *t* test. (C,D) HCT116 cells were treated with DMSO and increasing doses of NVP-2 and iCDK9 (green) for 24 hr prior to preparation of whole cell extracts and detection of the indicated proteins by Western blotting.

**Supplementary Figure 6.**
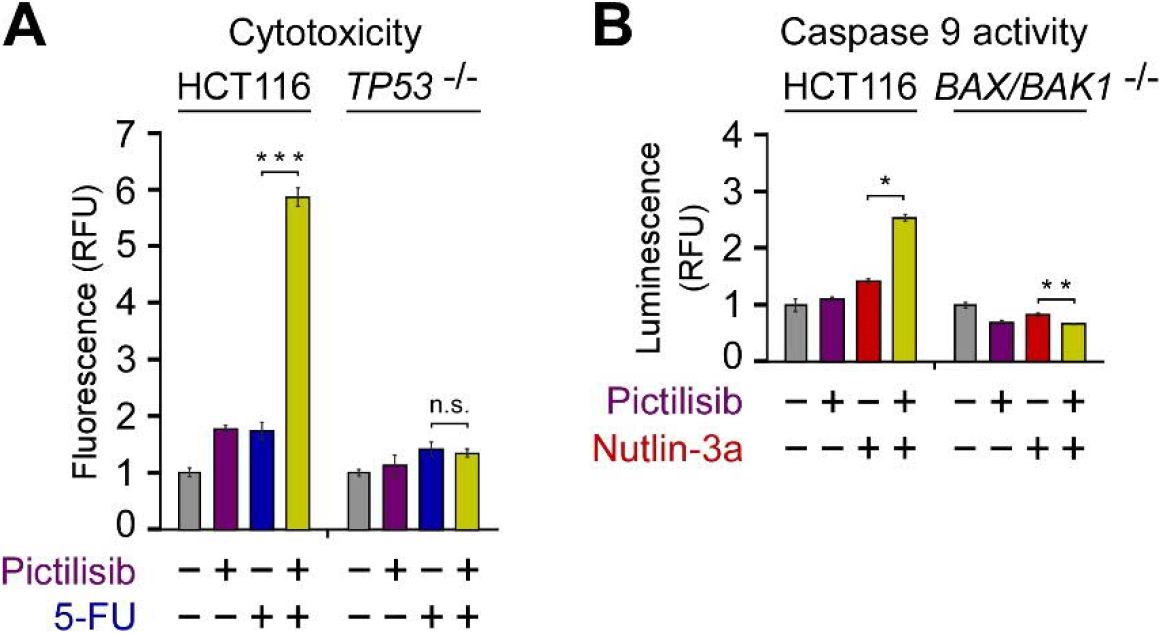
Repression of P-TEFb-dependent genes encoding key components of the pro-survival PI3K-AKT pathway is a driver of the synthetic lethality of p53 activation and P-TEFb inhibition. (A) Cytotoxicity of HCT116 and HCT116 *TP53* ^-/-^ cells treated with DMSO (grey), Pictilisib (1 μM; magenta), and 5-Fluorouracil (5-FU; 25 μM) (blue) alone and in combination (gold) as indicated for 48 hr measured using CellTox Green Cytotoxicity Assay. Results are presented as fluorescence values relative to the values of DMSO-treated cells and plotted as the mean ± s.e.m. (n = 3). ***, P < 0.001; n.s., non-significant, determined by Student’s *t* test. (B) Activity of Caspase 9 measured using Caspase-Glo 9 Assay in whole cell extracts of HCT116 and HCT116 *BAX*/*BAK1* ^-/-^ cells treated with DMSO (grey), Pictilisib (1 μM; magenta), and Nutlin-3a (10 μM; red) alone and in combination (gold) as indicated for 18 hr. Results are presented as luminescence values relative to the values of DMSO-treated cells and plotted as the mean ± s.e.m. (n = 3). *, P < 0.05; **, P < 0.01; n.s., non-significant, determined by Student’s *t* test.

**Supplementary Figure 7.**
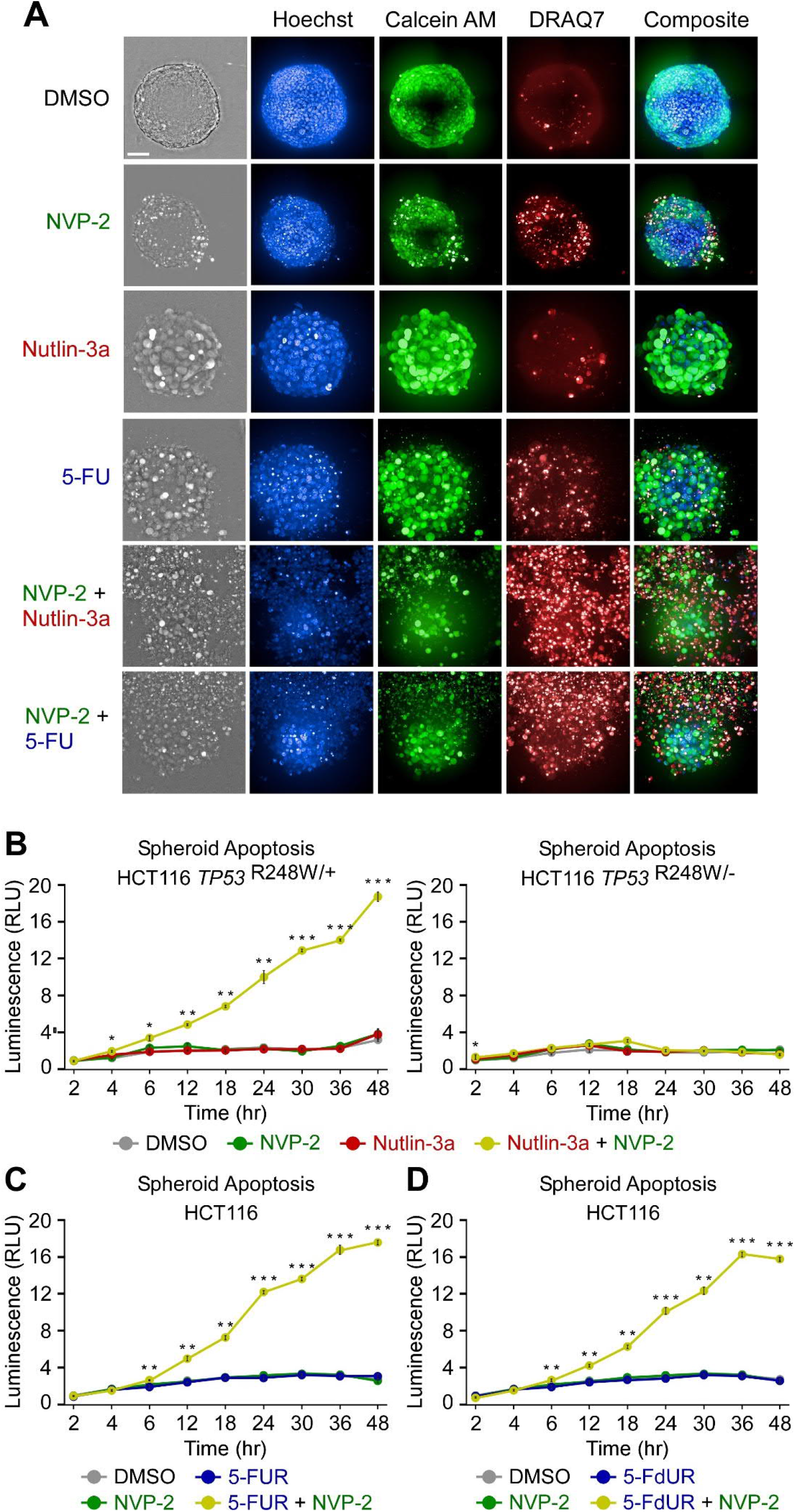
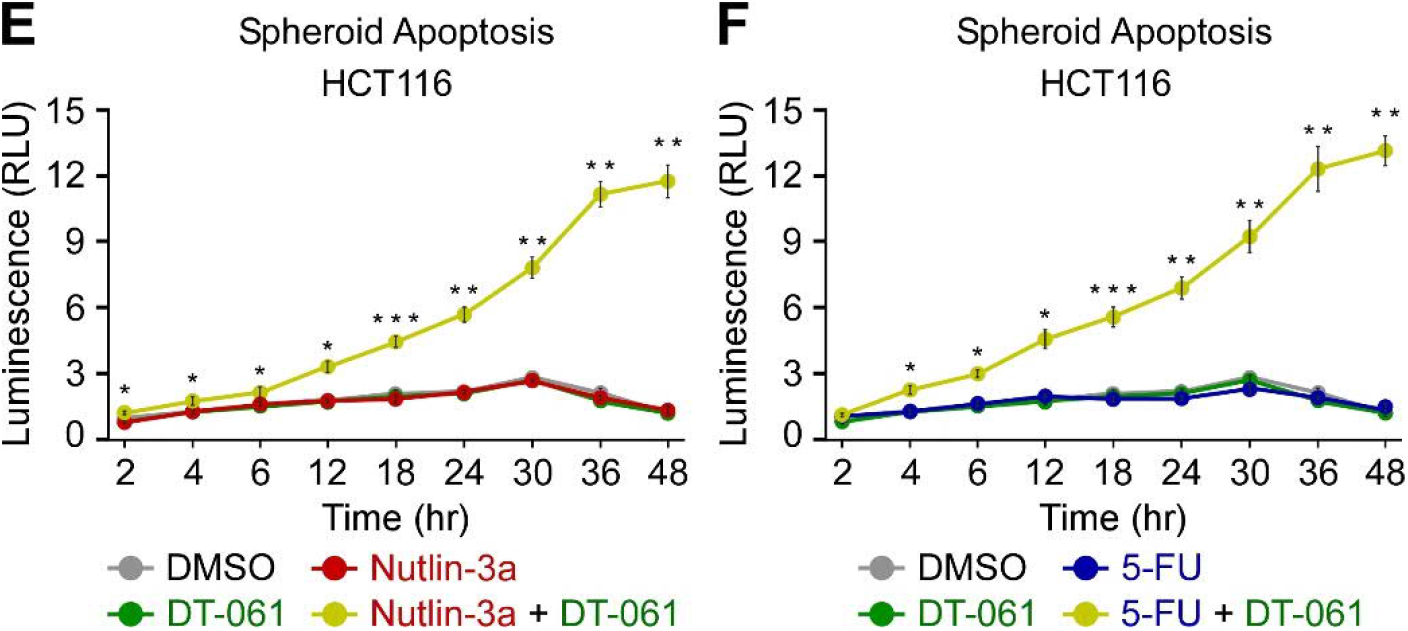
Combination treatments within the framework of p53 activation and P-TEFb inhibition trigger apoptosis of HCT116 spheroid cultures. (A) Representative images of HCT116 spheroid cultures treated with DMSO, Nutlin-3a (10 μM), NVP-2 (10 nM), and 5-Fluorouracil (5-FU; 25 μM) alone and in the combinations as indicated. Spheroids were formed for 48 hr, exposed to the treatments for 96 hr and stained with a mixture Calcein AM (1 μM), DRAQ7 (6 μM) and Hoechst (1x) prior to confocal microscopy imaging. Composite images of all three staining are shown on the right. Scale bar, 100 μm. (B-F) Apoptosis of the indicated HCT116 *TP53* ^R248W/+^, HCT116 *TP53* ^R248W/-^ and HCT116 cell spheroid cultures treated with DMSO (grey), Nutlin-3a (10 μM; red), NVP-2 (10 nM; green), 5-fluorouridine (5-FUR; 0.5 μM) and 5-fluorodeoxyuridine (5-FdUR; 0.15 μM), 5-Fluorouracil (5-FU; 25 μM) and DT-061 (10 μM; green) alone and in the indicated combinations (gold). Spheroids were formed for 48 hr prior to the treatments. Results obtained at the time points indicated below the graphs using RealTime-Glo Annexin V Apoptosis and Necrosis Assay are presented as luminescence values relative to the levels of DMSO-treated cells at 2 hr and plotted as the mean ± s.e.m. (n = 3). *, P < 0.05; **, P < 0.01; ***, P < 0.001, determined by Student’s *t* test using Nutlin-3a and Nutlin-3a + NVP-2 (B), 5-FUR and 5-FUR + NVP-2 (C), 5-FdUR and 5-FdUR + NVP-2 (D), Nutlin-3a and Nutlin-3a + DT-061 (E), and 5-FU and 5-FU + DT-061 (F) data sets.

**Supplementary Table 1a.** Antibodies used in the study.

| Antibody | Source | Identifier |
| --- | --- | --- |
| Mouse monoclonal anti-CDK9 | Santa Cruz Biotechnology | Cat#sc-13130; RRID: AB_627245 |
| Mouse monoclonal anti-p53 | Santa Cruz Biotechnology | Cat#sc-126; RRID: AB_628082 |
| Rabbit polyclonal anti-Cleaved PARP | Cell Signaling Technology | Cat#9541; RRID: AB_331426 |
| Mouse monoclonal anti-p21 | Santa Cruz Biotechnology | Cat#sc-53870; RRID: AB_785026 |
| Mouse monoclonal anti-GAPDH | Santa Cruz Biotechnology | Cat#sc-32233; RRID: AB_627679 |
| Rabbit polyclonal anti-RNA polymerase II CTD repeat YSPTSPS (phospho S2) | Abcam | Cat#ab5095; RRID: AB_304749 |
| Rabbit polyclonal anti-RNA polymerase II CTD repeat YSPTSPS (phospho S5) | Abcam | Cat#ab5131; RRID: AB_449369 |
| Rabbit monoclonal anti-RNA Polymerase II RPB1 NTD | Cell Signaling Technology | Cat#14958S; RRID: AB_2687876 |
| Normal rabbit IgG | Santa Cruz Biotechnology | Cat#sc-2027; RRID: AB_737197 |
| Mouse monoclonal anti-RNA Polymerase II RPB1 NTD | Santa Cruz Biotechnology | Cat#sc-17798; RRID: AB_677355 |
| Mouse monoclonal anti-p110a | Santa Cruz Biotechnology | Cat#sc-293172; RRID: AB_2847954 |
| Mouse monoclonal anti-p110β | Santa Cruz Biotechnology | Cat#sc-376641; RRID: AB_11150840 |
| Mouse monoclonal anti-p85a | Santa Cruz Biotechnology | Cat#sc-1637; RRID: AB_628126 |
| Anti-IRS-2 | Santa Cruz Biotechnology | Cat#sc-390761 |
| Mouse monoclonal anti-pan-Akt | Santa Cruz Biotechnology | Cat#sc-81434; RRID: AB_1118808 |
| Rabbit monoclonal anti-phospho-Akt (Ser473) | Cell Signaling Technology | Cat#4060; RRID: AB_2315049 |
| Rabbit monoclonal anti-phospho-p44/42 MAPK (Erk1/2) (Thr202/Tyr204) | Cell Signaling Technology | Cat#4376; RRID: AB_331772 |
| Rabbit monoclonal anti-p44/42 MAPK (Erk1/2) | Cell Signaling Technology | Cat#4695; RRID: AB_390779 |

**Supplementary Table 1b.** Chemicals used in the study.

| Chemical | Source | Identifier |
| --- | --- | --- |
| FO5A oncology library | FIMM | N/A |
| NVP-2 | MedChemExpress | Cat#HY-12214A |
| THAL-SNS-032 | Nathanael S. Gray Laboratory (Stanford University) | N/A |
| i-CDK9 | Qiang Zhou Laboratory (UC Berkeley) | N/A |
| Nutlin-3a | MedChemExpress | Cat#HY-10029 |
| 5-fluorouracil | Selleck Chemicals | Cat#S1209 |
| 5-fluorouridine | MedChemExpress | Cat#HY-107856 |
| 5-fluorodeoxyuridine | MedChemExpress | Cat#HY-B0097 |
| YKL-5-124 | Nathanael S. Gray Laboratory (Stanford University) | N/A |
| Senexin A | Selleck Chemicals | Cat#S8520 |
| OTS964 | Selleck Chemicals | Cat#S7648 |
| THZ531 | MedChemExpress | Cat#HY-103618 |
| Triptolide | Selleck Chemicals | Cat#S3604 |
| Pictilisib | MedChemExpress | Cat#HY-50094 |
| DT-061 | Jukka Westermarck Laboratory (University of Turku) | N/A |
| EDTA-free Protease Inhibitor Cocktail | Sigma | Cat#11873580001 |
| Random hexamers | Thermo Fisher Scientific | Cat#N8080127 |
| TRI Reagent | Sigma | Cat#T9424 |
| Calcein AM | Invitrogen | Cat#L3224A |
| DRAQ7 | Invitrogen | Cat#D15106 |
| Hoechst 33342 | Invitrogen | Cat#R37605 |

**Supplementary Table 1c.** Commercial assays used in the study.

| **Assay** | **Source** | **Identifier** |
| --- | --- | --- |
| CellTox™ Green Cytotoxicity Assay | Promega | Cat#G8731 |
| CellTiter-Glo® 2.0 Cell Viability Assay | Promega | Cat#G242 |
| RealTime-Glo Annexin V Apoptosis and Necrosis Assay | Promega | Cat#JA1011 |
| Caspase-Glo® 9 Assay | Promega | Cat#G8210 |
| RNeasy Mini Kit (50) | Qiagen | Cat#74104 |
| NEBNext Ultra Directional RNA Library Prep Kit | New England Biolabs | Cat#E7420 |
| NEBNext Poly(A) mRNA Magnetic Isolation Module | New England Biolabs | Cat#E7490 |
| M-MLV reverse transcriptase | Thermo Fisher Scientific | Cat#28025-013 |
| Turbo DNA-free™ kit | Thermo Fisher Scientific | Cat#AM1907 |
| Dynabeads Protein G | Thermo Fisher Scientific | Cat#10004D |
| FastStart Universal SYBR Green QPCR Master (Rox) | Sigma | Cat#4913914001 |
| MycoplasmaCheck detection kit | Eurofins | Cat#50400400 |

**Supplementary Table 1d.** Cell lines used in the study.

| Cell line | Source | Identifier |
| --- | --- | --- |
| HCT116 and HCT116 TP53<sup>-/-</sup> | Joaquin M. Espinosa<br>Laboratory (University of<br>Colorado) | N/A |
| HCT116 TP53<sup>R248W/+</sup> and HCT116 TP53<sup>R248W/-</sup> | Bert Vogelstein<br>Laboratory (John<br>Hopkins University) | N/A |
| HCT116 BAX/BAK1<sup>-/-</sup> | Ana J. Garcia-Saez<br>Laboratory (University of<br>Cologne) | N/A |
| CCD 841 CoN | ATCC | Cat#CRL-1790 |

**Supplementary Table 1e.** Software and algorithms used in the study.

| Software | Source | Identifier |
| --- | --- | --- |
| FastQC | Babraham Bioinformatics | http://www.bioinformatics.babraham.ac.uk/projects/fastqc/ |
| ContextMap v2.7.9 | (Bonfert et al., 2015)<sup>68</sup> | https://www.bio.ifi.lmu.de/software/contextmap/index.html |
| featureCounts | (Liao et al., 2014)<sup>70</sup> | http://bioinf.wehi.edu.au/featureCounts/ |
| edgeR | (Robinson et al., 2010)<sup>72</sup> | https://bioconductor.org/packages/release/bioc/html/edgeR.html |
| Molecular Signatures Database v6.0 | GSEA - Broad Institute (Subramanian et al., 2005)<sup>77</sup> | http://software.broadinstitute.org/gsea/msigdb/index.jsp |
| Harmony v4.9 High-Content Imaging and Analysis Software | PerkinElmer | N/A |
| MxPro QPCR Software v4.10 | Stratagene | N/A |

**Supplementary Table 2.**
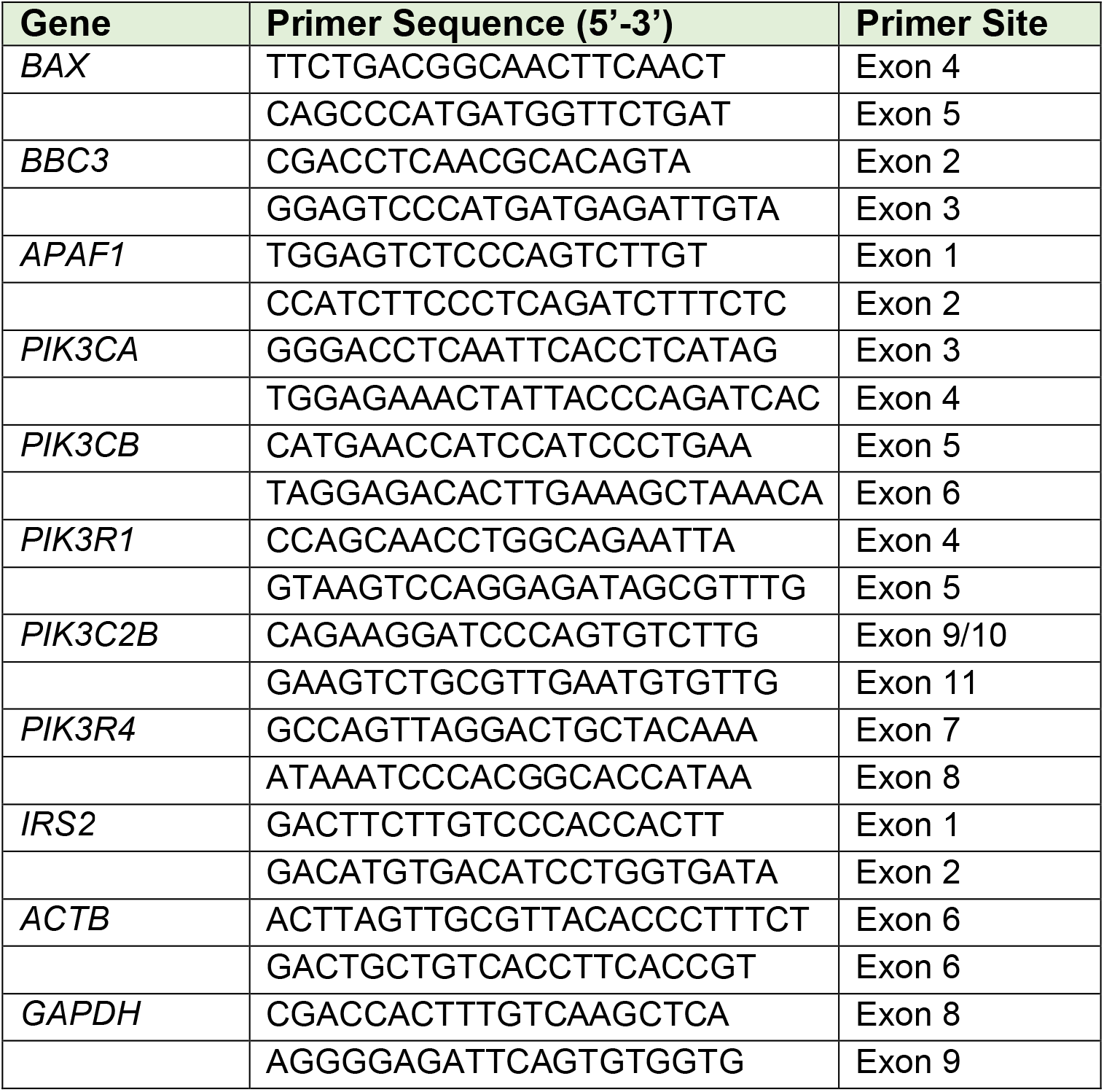
DNA oligonucleotides used in RT-qPCR assay.

**Supplementary Table 3.**
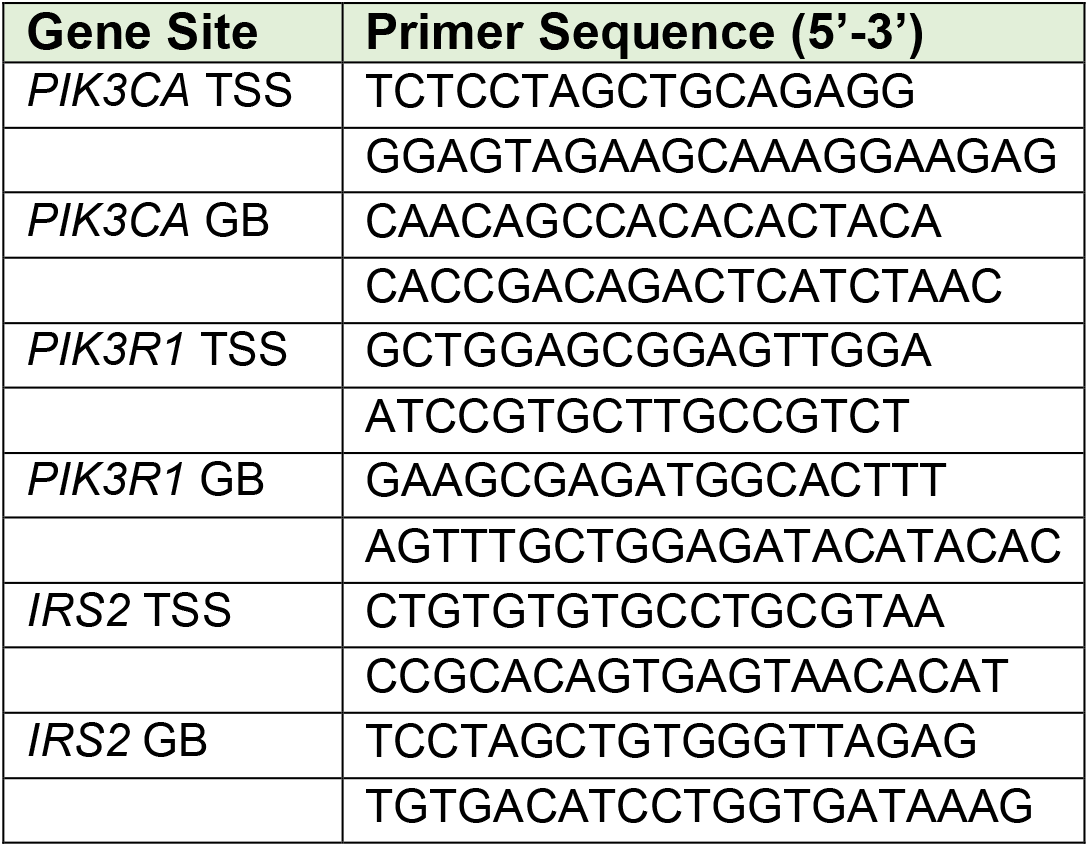
DNA oligonucleotides used in ChIP-qPCR assay.

